# Toward Robust Characterization of Dynamic Binding Pockets: Lessons from the HBV Capsid Assembly Modulator Site

**DOI:** 10.64898/2026.08.06.743403

**Authors:** Carolina Pérez-Segura, Liam W. Scott, Adam Zlotnick, Jodi A. Hadden-Perilla

**Affiliations:** Department of Chemistry & Biochemistry, University of Delaware, Newark, DE; Department of Molecular & Cellular Biochemistry, Indiana University, Bloomington, IN

## Abstract

Protein function often depends on ligand binding pockets that fluctuate among conformational states, altering their size, shape, topology, and accessibility, yet quantitative comparison of these dynamic cavities remains challenging because their boundaries are often inherently ambiguous. The *measure volinterior* algorithm uses fuzzy-boundary detection to characterize enclosed molecular spaces; here, the hepatitis B virus (HBV) capsid assembly modulator (CAM) binding site is used as a model system to develop and validate a practical workflow for applying the method to dynamic protein binding pockets. The resulting methodology provides practical guidance for parameter selection and evaluation, establishes a standardized protocol for quantitative characterization of the HBV CAM pocket, and demonstrates robust, reproducible performance across conformational ensembles derived from molecular dynamics (MD) simulations. More broadly, this work provides a reproducible strategy for adapting *measure volinterior* to other dynamic binding pockets, enabling consistent comparison of pocket geometry among independent structural studies.

**Graphical Abstract:** 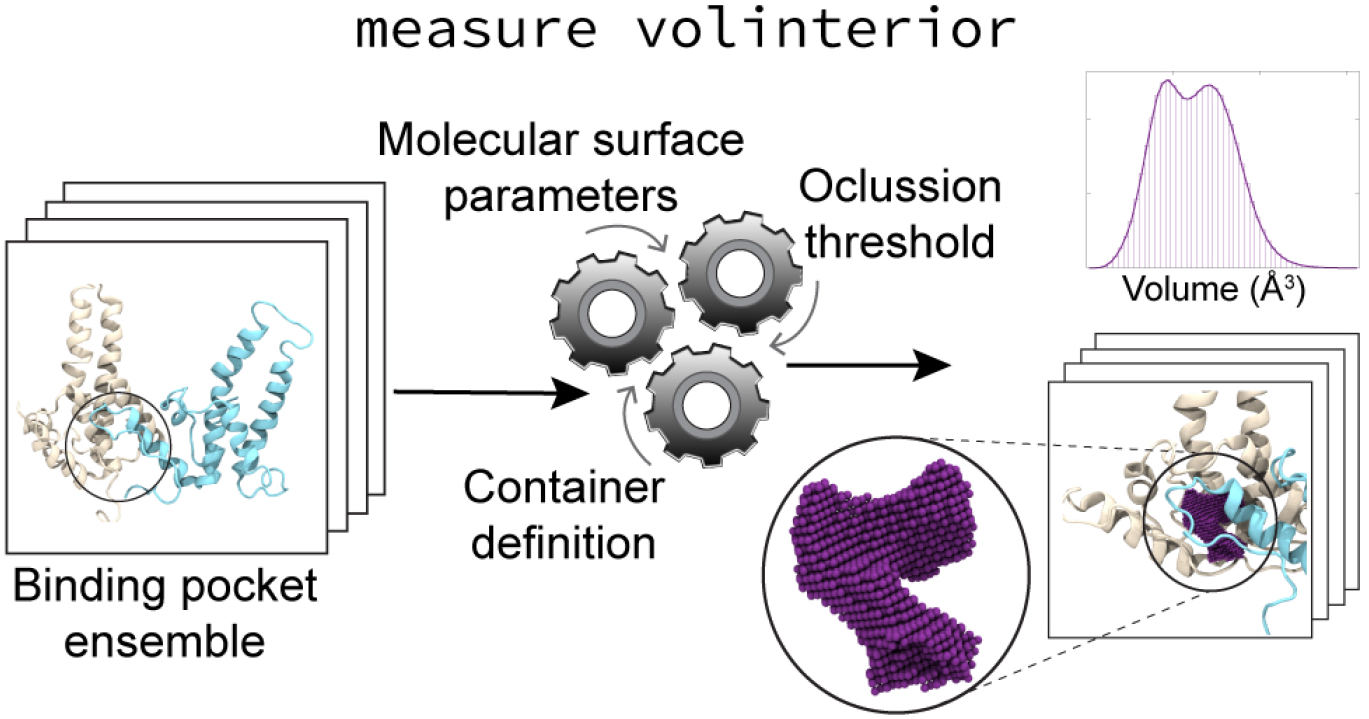

## Introduction

Characterization of binding pockets in proteins is central to structural biology and structure-based drug design, where pocket size, shape, and topology influence ligand recognition, affinity, and specificity. [1–4]. Binding pockets, however, are not static structural entities. Rather, proteins fluctuate among ensembles of conformations that alter pocket geometry and accessibility. Consequently, defining the boundary of a binding pocket becomes inherently ambiguous, particularly when comparing apo and ligand-bound structures, independent experimental models, and diverse conformations extracted from molecular dynamics (MD) simulations [5]. These challenges are especially pronounced for pockets that are partially open, flexible, or formed at protein–protein interfaces, where no unique physical boundary separates the cavity from the surrounding solvent environment.

A wide range of computational approaches has been developed to characterize binding pockets, including grid-based [6–9], probe-based [10–12], rolling-ball-based [13, 14], surface-based [15], and Voronoi-based [1, 16–18] methods. Although these approaches differ substantially in implementation, they share a common challenge: quantitative descriptions of pocket geometry depend critically on how the molecular boundary is defined. Differences in boundary definition can alter calculated pocket volumes, shapes, and topologies, complicating comparisons across independent studies and across structural ensembles generated by MD simulations. Furthermore, parameterizations that perform well for a single experimental structure may not remain suitable across the broader conformational landscape sampled by dynamic proteins or provide the computational efficiency required for high-throughput analysis of large structural ensembles.

The *measure volinterior* method [19], implemented in Visual Molecular Dynamics (VMD) [20], provides an attractive framework for characterization of dynamic binding pockets because it employs a tunable fuzzy-boundary definition of enclosed space, does not require prior knowledge of a bound ligand, and scales efficiently to large conformational datasets. The method was originally developed to characterize large molecular containers such as virus capsids, vesicles, and envelopes [21–26], where enclosed volumes are naturally delineated by a surrounding molecular shell. More recently, *measure volinterior* has also been applied to binding pockets [27–29], demonstrating its ability to quantify changes in pocket volume, geometry, and topology associated with ligand binding and conformational change.

Although the underlying *measure volinterior* algorithm is equally applicable to large molecular containers and small binding pockets, application to protein cavities introduces additional challenges. Because binding pockets occupy comparatively small volumes, relatively modest changes in molecular surface definition can produce large changes in the detected cavity geometry, topology, and calculated volume. Moreover, binding pockets frequently remain partially solvent accessible and may undergo substantial conformational fluctuations, requiring parameterizations that remain robust across diverse structural states while preserving a physically meaningful representation of the cavity. Practical guidance for developing, validating, and reproducing such parameterizations has not previously been established.

The hepatitis B virus (HBV) capsid assembly modulator (CAM) binding site provides an ideal test case for developing such a workflow because it combines substantial conformational heterogeneity with a dynamic protein–protein interface whose geometry is particularly sensitive to how cavity boundaries are defined. In its predominant *T* = 4 form, the HBV capsid is assembled from 120 core protein (Cp) dimers whose constituent chains adopt four quasi-equivalent conformations, denoted A, B, C, and D (**Figure 1A**); these chains are grouped into AB and CD dimers. Contact of these quasi-equivalent dimers within the capsid lattice creates structurally similar but non-identical interdimer interfaces containing hydrophobic binding pockets (**Figure 1B**). These quasi-equivalent pockets are formed by residues contributed from base and capping Cp chains and serve as the binding sites for CAMs, an important class of antiviral compounds under investigation for treatment of chronic HBV infection [30, 31]. Previous experimental and computational studies have shown that the CAM binding pocket exhibits substantial variability arising from quasi-equivalence, ligand occupancy, and conformational dynamics [29, 32–34], making it a particularly demanding system for evaluating cavity characterization methods.

**Figure 1:**
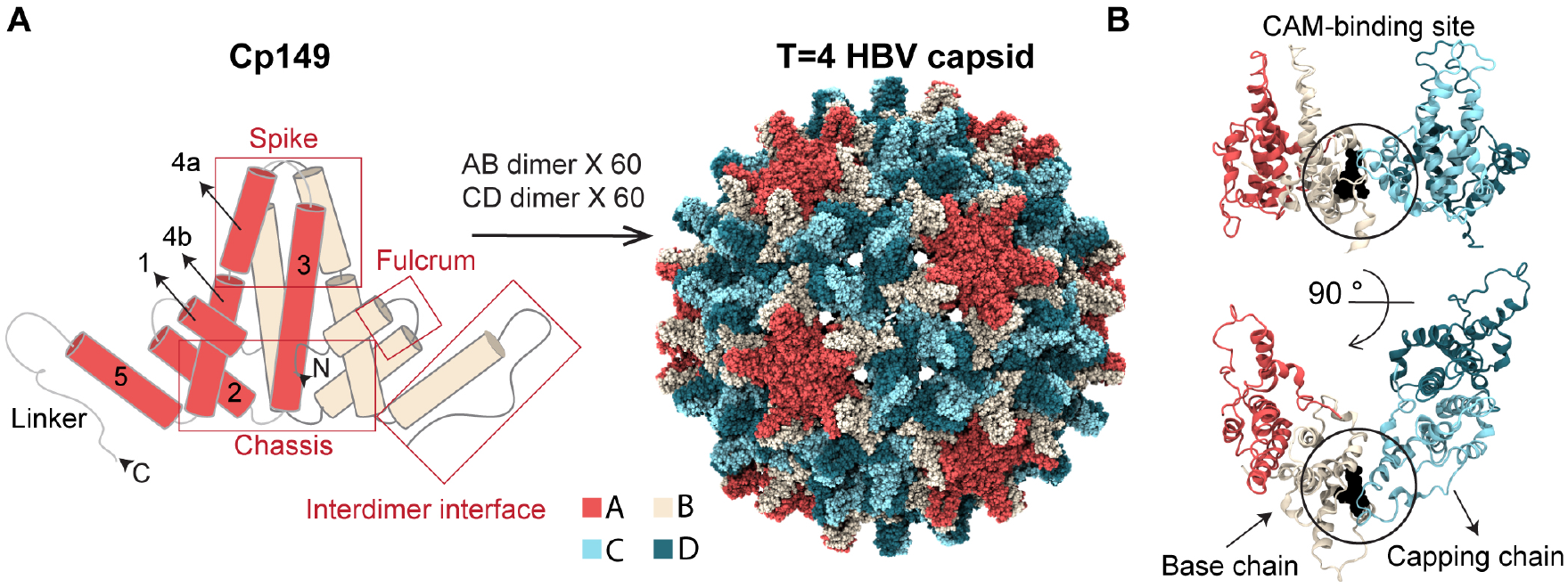
Architecture of HBV Cp, *T* = 4 capsid, and CAM binding site. (**A**) Schematic of the HBV core protein (Cp) assembly domain homodimer (residues 1-149, Cp149), indicating the five *α*-helices, the linker, and the structural subdomains defined by the tertiary and quaternary organization of Cp: chassis, fulcrum, spike, and the helix–loop motif that participates in the interdimer interface. The Cp homodimer can occupy two different quasi-equivalent positions in the *T* = 4 HBV capsid, denoted AB and CD, which are unique in their conformation and neighbor contacts. The four resulting chain types are colored here as A (red), B (beige), C (cyan), and D (blue). (**B**) The Cp interdimer interface is formed between a base chain from one homodimer and a capping chain from an adjacent homodimer, where the helix–loop motif of the capping chain packs against that of the base chain. Together, these structural elements create a hydrophobic pocket at the interdimer interface that serves as the binding site for capsid assembly modulators (CAMs), a class of small molecules that alter HBV capsid formation and stability. Here, the quasi-equivalent B-site is shown.

Here, the HBV CAM site is used as a model system to establish a practical framework for applying *measure volinterior* to dynamic binding pockets. Three interdependent components governing pocket characterization with *measure volinterior* are systematically evaluated: (i) definition of the molecular container, (ii) construction of the molecular surface, and (iii) selection of the occlusion threshold defining the effective pocket boundary. Using the HBV CAM site as an illustrative example, we develop an iterative workflow for optimizing these coupled parameters, assess their reproducibility, and demonstrate strategies for validating parameter performance across conformational ensembles. In addition to establishing a standardized protocol for characterization of the HBV CAM site, this work provides practical guidance for developing robust and reproducible *measure volinterior* parameterizations for other dynamic binding pockets.

## Methods

### Computational environment and hardware requirements

The *measure volinterior* method [19] is available in VMD [20] version 1.9.3 or later. VMD is supported on Linux, Windows, and macOS operating systems and can be obtained as source code or precompiled binaries from the VMD website. Although *measure volinterior* can be executed using either CPU or GPU implementations, the GPU implementation should be considered a required component for reproducing the results described here. The CPU and GPU implementations do not produce identical pocket geometries or volume measurements, and all parameter development, optimization, and validation presented in this work were performed using the GPU implementation. Consequently, the recommended parameter ranges presented below are specific to GPU-generated results and should not be assumed to transfer directly to CPU calculations. Execution of the GPU implementation requires an NVIDIA graphics processing unit with CUDA support. Compatibility between VMD, the installed NVIDIA driver version, and the CUDA runtime environment is required for proper GPU detection and execution. Users should therefore verify compatibility among the VMD release, GPU drivers, and CUDA version prior to performing calculations. Additional information regarding CUDA support and GPU acceleration in VMD, including recommended GPU hardware and compatibility requirements, is available from the VMD documentation.

### Principles of pocket detection with *measure volinterior*

The *measure volinterior* method [19] is designed to identify and characterize enclosed spaces within biomolecular systems. Although originally developed for large molecular containers such as entire virus capsids and lipid vesicles, the method can also be applied to dynamic binding pockets given careful selection of the input parameters governing pocket definition. To identify the space associated with a binding pocket, *measure volinterior* uses a three-dimensional ray-casting approach to classify the empty space surrounding the molecular surface. Briefly, the system is discretized into a three-dimensional grid of volume elements, or voxels, and rays are projected outward from each voxel in different directions. The classification of a voxel as belonging to the pocket interior depends on whether rays cast from that voxel intersect the surrounding molecular surface. Consequently, the accuracy of pocket characterization depends strongly on how the molecular surface is represented.

Unlike completely enclosed molecular containers, binding pockets are open systems that typically remain partially solvent accessible. Application of *measure volinterior* to binding pockets therefore requires use of the fuzzy-boundary detection feature [19]. Rather than assigning voxels as strictly interior or exterior, fuzzy-boundary detection quantifies the degree to which each voxel is occluded by the surrounding molecular surface. Specifically, the number of rays intersecting the molecular surface is normalized by the total number of rays cast from each voxel to produce an occlusion score ranging from 0 to 1. Voxels with scores approaching unity are more deeply buried within the pocket, whereas voxels with lower scores remain more exposed to the solvent environment. Selection of an occlusion threshold subsequently defines the effective boundary of the pocket across its opening. This feature enables characterization of partially open binding pockets by allowing the enclosed space associated with the cavity to be delineated according to the surrounding protein architecture. As a result, *measure volinterior* can be used to characterize binding pockets even in highly dynamic systems.

### Interdependent parameters governing pocket definition

Application of *measure volinterior* requires specification of (i) a VMD atom-selection defining the molecular container, (ii) four parameters governing construction of the molecular surface and ray-casting calculations, and (iii) an occlusion threshold defining the effective boundary of the pocket across its opening. Importantly, these values are system dependent and must be chosen by the user to faithfully represent the structural features of the system of interest. Furthermore, because the container definition, molecular surface, and pocket boundary collectively determine the resulting cavity geometry, these parameters are coupled and cannot be optimized independently.

The first of the three interdependent parameterization components governing pocket characterization with *measure volinterior* is definition of the molecular container. In the present application, the molecular container is not the entire protein but rather the localized subset of residues that defines the binding pocket of interest. In VMD, this container is specified through an atom-selection, which determines which structural elements contribute to the molecular surface surrounding the detected cavity. Importantly, the entire biomolecular structure need not, and typically should not, be provided as input to *measure volinterior*. Because fuzzy-boundary detection classifies voxels according to their occlusion by the molecular surface, the method evaluates space surrounding the entire atom-selection, potentially detecting enclosed regions unrelated to the binding pocket. Users should therefore define a localized molecular container consisting only of residues that contribute to the pocket architecture. Selected residues need not comprise a continuous amino acid sequence, but complete residues are generally preferred over isolated atoms because they provide a more robust definition of the molecular surface that is less sensitive to conformational fluctuations. Restricting the atom-selection to the local structural environment surrounding the pocket not only improves computational efficiency, but also reduces inclusion of unrelated surface depressions, minimizes isolated voxels and disconnected cavity regions, and preserves a cleaner and more interpretable pocket boundary.

The second of the three interdependent parameterization components governing pocket characterization with *measure volinterior* is construction of the molecular surface. VMD’s QuickSurf representation [35] is used to generate the molecular surface that serves as the steric boundary interrogated by *measure volinterior*. Rays cast from each voxel are tested for intersection with this surface to determine the degree of occlusion. Consequently, the detected cavity is inherently sensitive to how the molecular surface is represented. QuickSurf is governed by three user-defined parameters: Radius Scale, Isovalue, and Grid Spacing. These parameters collectively determine the smoothness, compactness, and spatial resolution of the molecular surface and are therefore strongly coupled. Here, the default VMD van der Waals (vdW) radii were used for all atoms during surface construction. If needed, alternative atomic radii sets can be loaded from biomolecular force fields, implicit solvent models, or element definitions included in standard-format PDBs. Radius Scale (Å) controls scaling of the vdW radii used to generate the molecular surface and therefore influences the effective bulkiness of the representation. Increasing the Radius Scale expands the molecular surface outward from the atomic coordinates, whereas smaller values produce a more compact surface. Isovalue defines the density threshold used to generate the surface from overlapping atomic density distributions. Higher Isovalue values generally produce tighter surfaces that contour more closely around the atomic core density, whereas lower values generate smoother and more expanded surfaces. Grid Spacing (Å) defines the spatial resolution of the voxel grid used during surface construction. Smaller Grid Spacing values produce finer voxelization and greater structural detail but increase computational cost.

In addition to the parameters governing surface construction, *measure volinterior* requires specification of the Number of Rays to be cast from each voxel. Increasing the Number of Rays improves angular sampling of the surrounding molecular surface and therefore increases the accuracy with which occlusion is evaluated. However, larger Number of Rays also increases computational cost and provides diminishing improvements once the surrounding surface has been sufficiently sampled. Consequently, the Number of Rays should balance computational efficiency with adequate sampling of the local molecular environment.

The third of the three interdependent parameterization components governing pocket characterization with *measure volinterior* is selection of the occlusion threshold. Because binding pockets are typically partially solvent accessible rather than fully enclosed containers, cavity detection should be performed using fuzzy-boundary detection [19]. Under this approach, each voxel is assigned an occlusion score corresponding to the fraction of rays emitted from that voxel that intersect the surrounding molecular surface. Equivalently, this value may be interpreted as the probability that the voxel belongs to the cavity interior according to the ray-casting procedure. An occlusion threshold is subsequently applied to determine which voxels are retained as part of the detected cavity.

Operationally, this threshold defines the effective boundary of the pocket across its solvent-accessible opening. Higher thresholds restrict the detected cavity to more deeply buried regions of the pocket, whereas lower thresholds permit inclusion of increasingly solvent-exposed regions. When using fuzzy-boundary detection, the combination of the `-probmap` and `-count_pmap` flags returns voxel counts corresponding to successive occlusion-score thresholds ranging from 0.1 to 0.9 (i.e., 10% to 90% occlusion). For example, element 8 of the returned list corresponds to an occlusion threshold of 0.9 and therefore includes only voxels with occlusion scores greater than or equal to 0.9. Importantly, the occlusion threshold should not be interpreted as defining a uniquely correct physical boundary for the pocket. Rather, it provides a tunable definition of the cavity opening whose suitability depends on the structural features and conformational behavior of the system being analyzed.

### Strategies for visualizing the detected pocket geometry

In addition to voxel counts that quantify cavity volume, *measure volinterior* generates a three-dimensional volumetric map describing the geometry of the detected pocket. When fuzzy-boundary detection is enabled, each voxel in the map is assigned an occlusion score corresponding to the degree to which the voxel is enclosed by the surrounding molecular surface. Application of an occlusion threshold then defines the effective boundary of the pocket across its solvent-accessible opening and delineates the three-dimensional volume of space associated with the cavity. Visualization of the resulting cavity geometry is an important component of parameter selection because physically reasonable pocket volumes should also produce physically reasonable pocket shapes and boundaries. The most effective way to evaluate the suitability of *measure volinterior* parameters for characterizing a given pocket is to visualize the results they produce.

The detected cavity volume can be visualized directly using VMD’s Isosurface representation. In this approach, the volumetric map generated by *measure volinterior* is rendered using the desired occlusion threshold to define the pocket boundary. Importantly, when visualizing the volumetric map using VMD’s Isosurface representation, the selected Isovalue should be set equal to the occlusion threshold used during *measure volinterior* cavity characterization in order to accurately reproduce the detected pocket geometry. This Isovalue determines which voxels from the map are displayed and is distinct from the Isovalue parameter used by the QuickSurf representation to generate the molecular surface. The spatial resolution of the Isosurface representation is determined by the Grid Spacing parameter used during construction of the molecular surface and calculation of the volumetric map.

An alternative visualization strategy is to represent the detected cavity as a three-dimensional point cloud using a regularly spaced lattice of dummy atoms. A Tcl script for generating a dummy-atom lattice centered on the detected cavity is provided in **Code Snippet S1**. This script uses VMD’s Voltool package to extract the origin and dimensions of the volumetric map and then places dummy atoms at voxel centers throughout the volume according to the value of Grid Spacing. By loading the volumetric map into the resulting dummy-atom structure, atom-selection expressions such as vol0 > 0.9 can be used to visualize only those voxels satisfying a chosen occlusion threshold. The point cloud representation is particularly useful for evaluating cavity continuity, identifying isolated voxels, and examining how parameter choices influence the detected pocket geometry. Because dummy-atom point clouds have a smaller disk footprint than volumetric maps, this strategy also represents a more efficient way to store and process the three-dimensional data produced by *measure volinterior* when analyzing large conformational ensembles.

The original *measure volinterior* publication used pre-equilibrated water boxes to visualize detected interior volumes, an approach that performs well for large molecular containers such as virus capsids [19]. However, visualization using water molecules can introduce artifacts when examining small binding pockets. Because water molecules are not uniformly distributed in space, their positions do not necessarily coincide with voxel centers. The characteristic oxygen–oxygen separation distance in liquid water is approximately 2.8–2.9 Å under typical experimental conditions [36, 37], substantially larger than the Grid Spacing recommended for cavity detection in small pockets, and local water organization depends on molecular orientation and hydrogen-bonding interactions. Consequently, visualizations that rely on pre-equilibrated water boxes may incompletely represent the detected volume or introduce apparent irregularities that do not reflect the underlying *measure volinterior* results. While pre-equilibrated water boxes are adequate for displaying the interior space of large molecular containers, visualization using a regular dummy-atom lattice provides a more faithful representation of the detected pocket geometry for small cavities.

## Results and Discussion

### Optimized *measure volinterior* protocol for the HBV CAM site

The HBV CAM site is highly mobile [34, 38] and provides an excellent model system for developing and evaluating of *measure volinterior* parameters for dynamic binding pockets. Four experimentally determined HBV capsid structures spanning apo and ligand-bound states were used throughout this work: apo-form (PDB 2G33, 3.96 Å [39]), AT130-bound (PDB 4G93, 4.2 Å [40]), HAP1-bound (PDB 2G34, 5.0 Å [39]), and HAP18-bound (PDB 5D7Y, 3.9 Å [41]). Missing residues within the linker region of each quasi-equivalent chain were previously modeled to generate complete assembly domain (residues 1-149, Cp149) structures suitable for MD simulation [34, 42, 43]. Unless otherwise noted, parameter development and evaluation were performed using this full-length construct commonly employed for in vitro studies of HBV.

The final optimized *measure volinterior* parameters selected for characterization of the HBV CAM site are summarized in **Table 1**. **Figure 2** illustrates the pocket geometries and corresponding volume measurements obtained using these parameters for the four quasi-equivalent CAM sites across the four HBV capsid crystal structures. The corresponding Tcl script for the VMD workflow is provided in **Code Snippet 1**. Detected pocket volumes are visualized as three-dimensional point clouds occupying the interdimer interface, providing a direct representation of the cavity geometry identified by *measure volinterior*.

**Figure 2:**
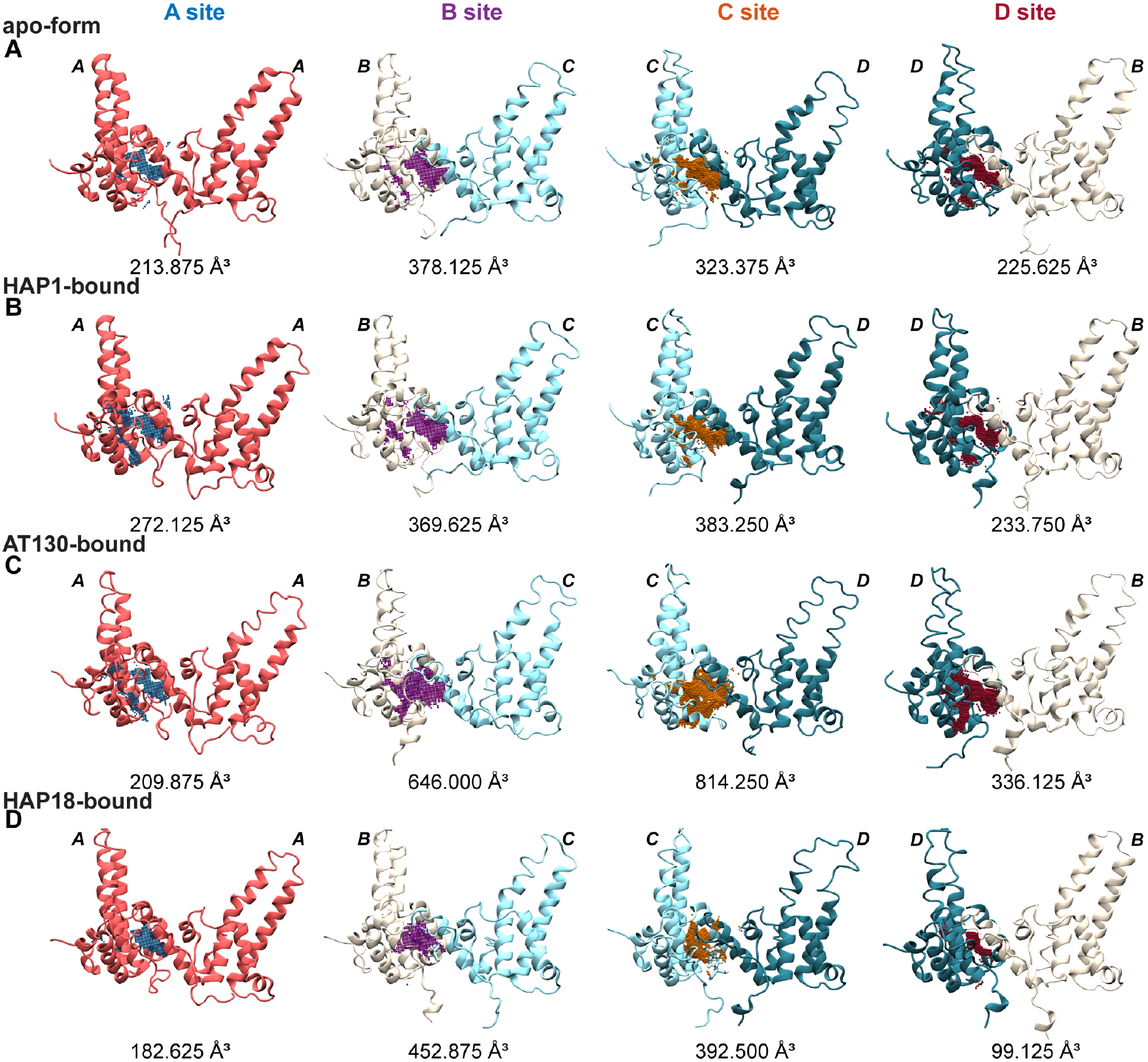
Quasi-equivalent CAM binding sites are different shapes and sizes. Pocket three-dimensional shape and volume results calculated by *measure volinterior* for the four quasi-equivalent CAM sites of four different HBV *T* = 4 capsid crystal structures: (**A**) apo-form (PDB 2G33 [39]), (**B**) HAP1-bound (PDB 2G34 [39]), (**C**) AT130-bound (PDB 4G93 [40]), and (**D**) HAP18-bound (PDB 5D7Y [41]). Volume calculations were performed using the optimized *measure volinterior* parameters summarized in Table 1. Pocket shapes represented as point clouds. Pocket volumes reported to three digits of precision to show that they are multiples of 0.125 Å^3^, the volume of a single voxel of Grid Spacing 0.5 Å.

**Table 1:**
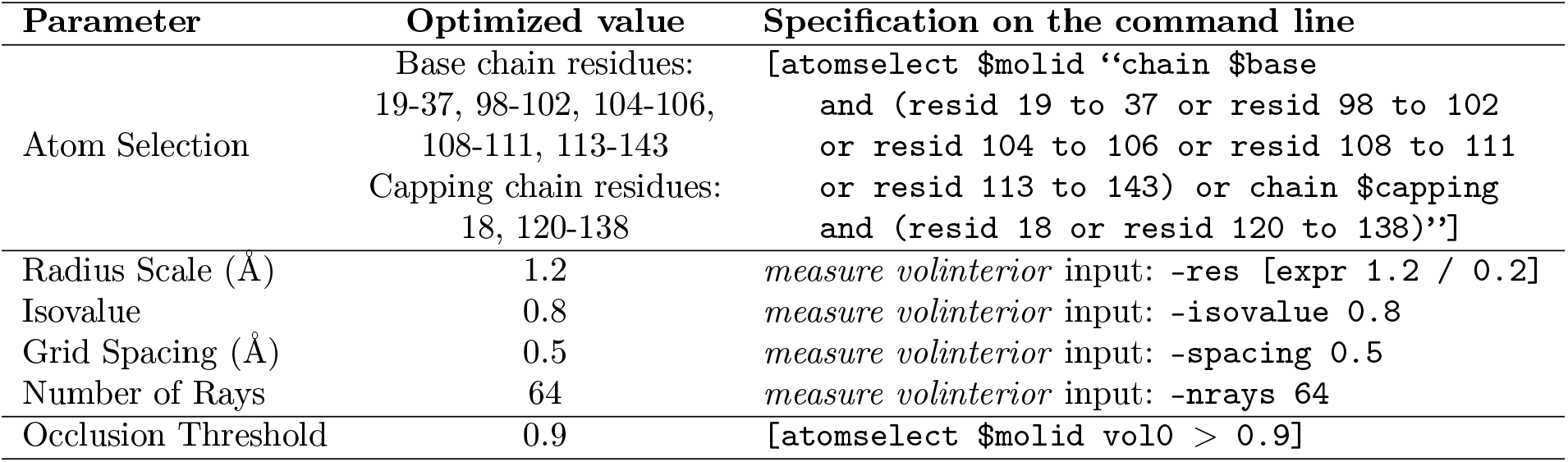
VMD *measure volinterior* parameters for the HBV CAM binding site.

Pronounced differences in pocket volume are observed even in the apo-form capsid, where all four quasi-equivalent CAM sites are unoccupied (**Figure 2A**). According to the crystal structure, pocket volumes range from 213.875 Å^3^ at the A-site to 378.750 Å^3^ at the B-site; that is, the B-site is 177% the size of the A-site. Thus, quasi-equivalence and local structural environment within the capsid lattice can substantially influence pocket geometry independent of ligand binding. Interestingly, the larger B- and C-sites correspond to the pockets occupied by CAMs in the ligand-bound crystal structures (**Figure 2B–D**), suggesting that quasi-equivalence may contribute to differences in ligand accessibility among the four interdimer interfaces. Despite being comparatively small, the volumes of the unoccupied A- and D-sites vary across the four crystal structures without exhibiting a clear dependence on CAM chemotype (**Figure 2B–D**), highlighting the inherent structural variability of the CAM binding pocket.

The optimized *measure volinterior* parameters summarized in **Table 1** were developed specifically for characterization of the HBV CAM site and validated using extensive atomistic MD simulations sampling millions of binding pocket conformations [34, 42, 43]. Unlike parameterizations based solely on one or a few experimental structures, this approach captures the broad conformational heterogeneity exhibited by the CAM site in intact HBV capsids and therefore provides a more stringent assessment of parameter robustness. Consequently, the selected parameters were optimized not simply to perform well for a single pocket conformation, but to provide consistent and physically meaningful characterization across the dynamic conformational landscape of the CAM site.

When applied using the GPU implementation of *measure volinterior* together with VMD’s default vdW radii, these optimized parameters produce reproducible cavity geometries and numerically identical volume measurements for identical input structures with identical three-dimensional coordinates. An example dimer-of-dimers encompassing the quasi-equivalent B-site CAM pocket extracted from PDB 2G33 [39], with the unresolved linker residues remodeled to generate the full-length Cp149 structure, is provided in the Supporting Information. Analysis of this structure using the workflow described in **Code Snippet 1** yields a cavity volume of 378.125 Å^3^, providing users with a straightforward reproducibility benchmark for validating correct implementation of the protocol. Importantly, this and all other HBV Cp models analyzed in this work were protonated prior to pocket characterization. Because experimental structures typically lack hydrogen atoms, their omission contracts the molecular surface surrounding the protein and can lead to overestimation of the binding pocket volume.

The optimized parameter set presented here provides a standard for characterization of the HBV CAM site and enables direct, reproducible comparison of pocket volume, geometry, and topology across independent structural studies. Because these parameters were developed specifically for the HBV CAM site, they should not be expected to transfer directly to unrelated binding pockets. Instead, the HBV CAM site serves here as a representative case study illustrating the iterative workflow used to develop, optimize, and validate a robust *measure volinterior* parameterization. The following sections describe this workflow and provide practical guidance for adapting the approach to other dynamic binding pockets.

#### Code Snippet 1: Tcl script to calculate CAM binding site volume with *measure volinterior*.

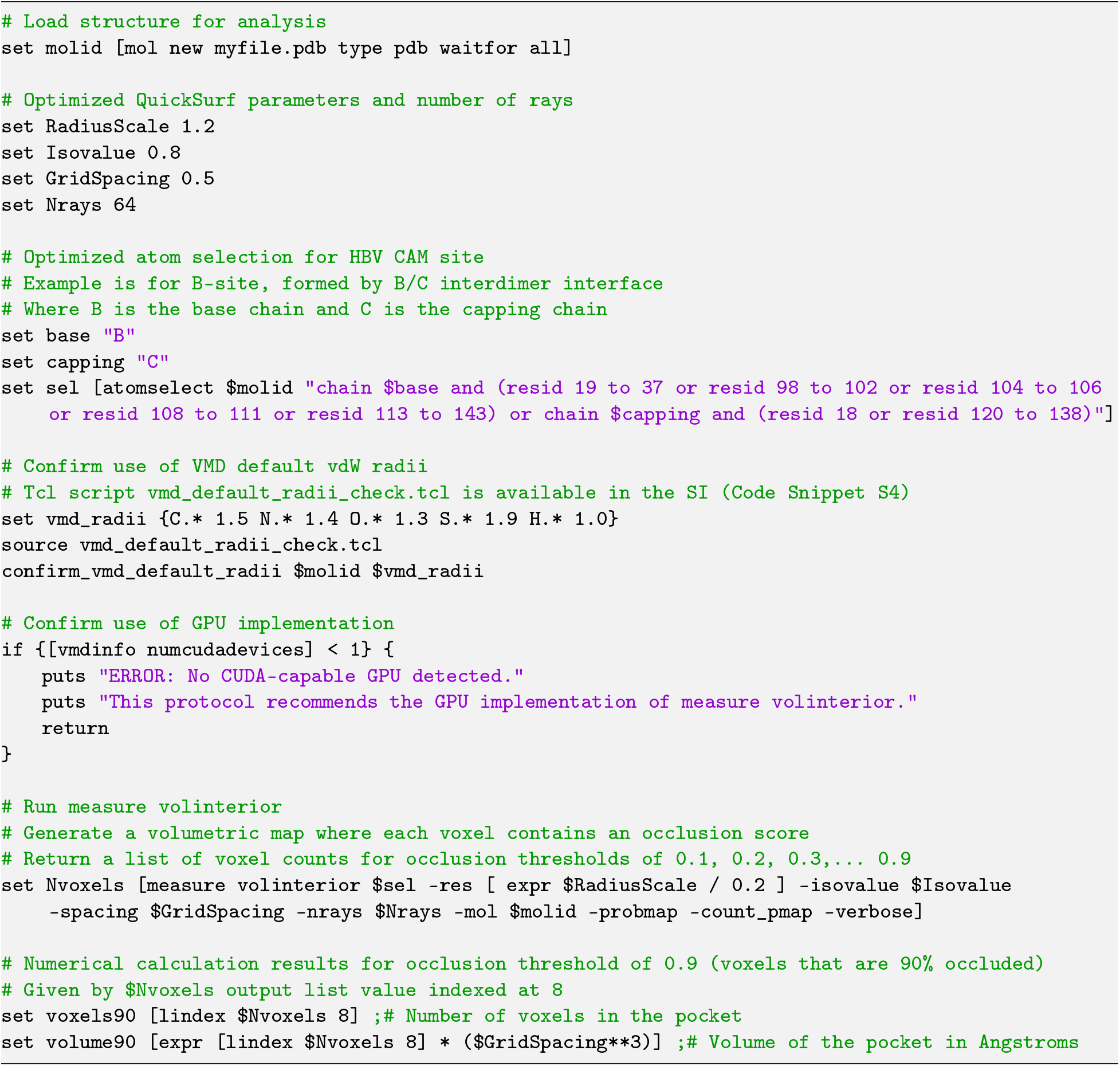

### Defining the binding pocket: Localizing the VMD atom-selection

The first step in applying *measure volinterior* to a binding pocket is defining the cavity through an appropriate VMD atom-selection. For the HBV CAM site, the goal is to construct a localized molecular container that captures the geometry of the binding pocket while excluding unrelated structural features that may confound cavity detection. Because the CAM site is formed at the interface between two neighboring Cp dimers, only the two monomer chains that directly contribute to the pocket are required (as shown in **Figure 2**). For example, characterization of a B-site pocket requires only the B (base) and C (capping) chains that form the corresponding interdimer interface.

Even after restricting the analysis to the two chains forming the pocket, inclusion of complete monomers introduces additional enclosed regions unrelated to the CAM site (**Figure 3A**). In addition to the CAM binding pocket, each HBV Cp contains a lipid binding pocket [28, 44, 45] The Cp surface also contains numerous other shallow depressions arising from tertiary structural organization and local side-chain packing. Because *measure volinterior* evaluates all space surrounding the selected molecular surface, these additional features are detected alongside the CAM site, producing substantial noise and complicating interpretation of the resulting cavity geometry. Consequently, further localization of the atom-selection is required to isolate the CAM binding pocket as the molecular container of interest.

**Figure 3:**
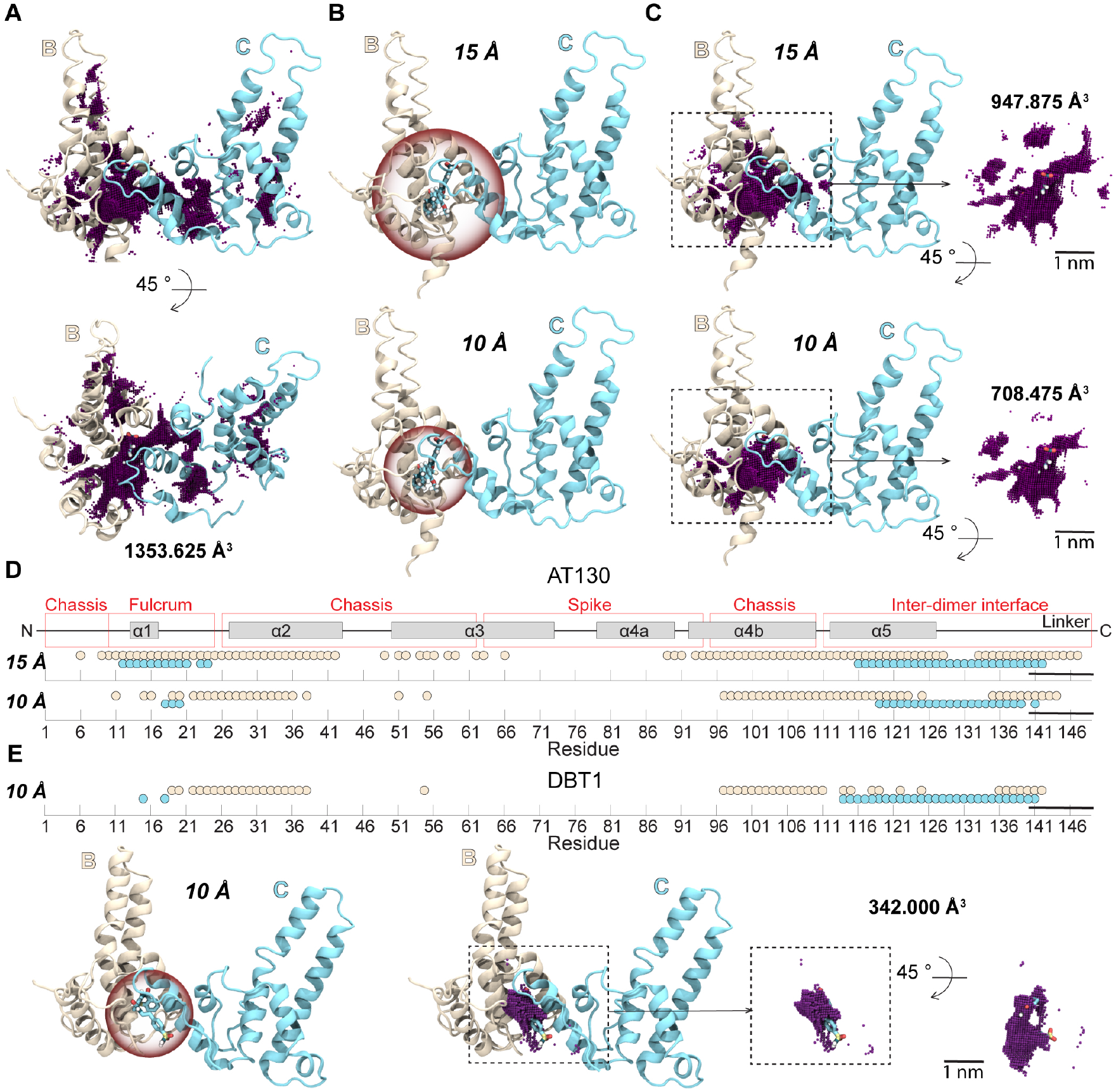
Ligand-guided definition of the CAM site. (**A**) Providing the complete Cp base and capping chains as the input atom-selection causes highly occluded regions throughout the protein structure to be detected by *measure volinterior*, including numerous cavities unrelated to the CAM site. (**B**) A ligand-guided definition localizes the molecular container to the binding site by selecting residues within a specified distance of a bound CAM (e.g., a 10–15 Å cutoff). (**C**) The resulting pocket geometry depends strongly on the selected distance cutoff. A shorter cutoff may omit residues that contribute to the cavity boundary but do not directly contact the ligand, whereas a longer cutoff incorporates unrelated enclosed regions outside the binding pocket. Even modest changes from a 10 Å to 15 Å cutoff produce substantially different cavity geometries. (**D**) The differences in detected pocket geometry arise from the distinct residue sets selected by each cutoff. The residues captured using 10 Å and 15 Å cutoffs around AT130 bound in the quasi-equivalent B-site (PDB 4G93 [40]) are compared. (**E**) Ligand-guided residue selection also depends on ligand identity. Residues selected using the same 10 Å cutoff differ substantially for AT130 and DBT1 (PDB 6WFS [46]) because the two CAMs adopt distinct binding modes within the same binding pocket. Consequently, a ligand-guided molecular container depends not only on the distance cutoff but also on how a particular ligand occupies the cavity. Volume calculations were performed using the optimized *measure volinterior* parameters summarized in Table 1.

A natural starting point is the extensive body of structural and biochemical data identifying residues important for CAM binding. Experimental studies have implicated residues including Phe23, Pro25, Thr33, Trp102, Ile105, Ser106, Tyr118, and Leu140 in the Cp base chain, together with Val120, Val124, Thr128, Arg133, and Pro134 in the capping chain, as key contributors to CAM recognition [32, 41, 47–50]. However, these residues alone do not fully define the geometry of the binding pocket. Rather, they identify amino acids that interact directly with the ligand, whereas cavity detection requires defining the broader molecular surface that encloses the pocket.

This distinction is particularly important for dynamic systems. MD simulations of the HBV capsid have demonstrated that interdimer interfaces undergo substantial conformational fluctuations and can form transient contacts not apparent in static experimental structures [34, 38]. Some of these dynamically sampled interactions have been validated experimentally as important determinants of capsid assembly and native CAM resistance [38]. Consequently, defining a molecular container solely from residues identified in a single, rigid experimental structure risks omitting structural features that contribute to pocket geometry elsewhere within the conformational ensemble. Whenever possible, the atom-selection used for *measure volinterior* should therefore be informed by a consensus view of the binding site derived from multiple experimental structures or conformational states rather than a single static model. In this regard, MD simulations can be particularly valuable because they sample the continuous conformational fluctuations of a binding pocket and provide a more comprehensive description of the molecular environment that forms the cavity.

Two complementary strategies can be used to define the molecular container surrounding a small molecule binding pocket: a ligand-guided approach and a pocket-guided approach. The ligand-guided approach identifies container residues according to their proximity to a bound ligand, whereas the pocket-guided approach defines the molecular container directly from the surrounding protein architecture. Both approaches seek to identify the structural elements that enclose the cavity of the binding pocket, but they differ fundamentally in the information used to define those elements. Ligand-guided definitions naturally incorporate induced-fit conformational changes captured in protein–ligand complexes and therefore provide a straightforward strategy when a representative bound structure is available. However, because the molecular container definition is derived from the ligand within, the resulting atom-selection may depend on ligand size, binding mode, and the extent to which the ligand occupies the cavity. Furthermore, this strategy cannot be applied to apo structures or complexes for which the ligand coordinates are not resolved. In contrast, a pocket-guided approach defines the molecular container from the protein architecture itself, making it independent of ligand identity while remaining applicable to both apo and ligand-bound conformations.

### Defining the binding pocket: Ligand-guided approach

For protein–ligand complexes, a natural strategy for defining the molecular container is to select residues within a specified distance of the bound ligand. This ligand-guided approach was successfully employed in a recent application of *measure volinterior* to characterize interactions of the inhibitor pirmitegravir with HIV-1 integrase [27]. To evaluate suitability of the approach for dynamic pockets like the HBV CAM site, residues containing at least one atom within 10 Å or 15 Å of the AT130 molecule were selected from the B-site of the AT130-bound capsid (**Figure 3B**). Although both distance cutoffs identify the general region surrounding the CAM site, increasing the cutoff from 10 Å to 15 Å introduces additional enclosed regions that are clearly unrelated to the binding pocket. As shown in **Figure 3C**, the larger cutoff produces isolated voxels and disconnected cavity components located far from the bound AT130 molecule that do not represent structural features of the CAM site. These extraneous regions fluctuate throughout an MD simulation, further complicating interpretation of the detected cavity. Consequently, the molecular container no longer isolates the cavity of interest, causing *measure volinterior* to characterize multiple enclosed regions rather than the CAM binding pocket alone.

This reduced specificity of the molecular container is reflected directly in the residues included in the two distance cutoffs. As shown in **Figure 3D**, the 10 Å and 15 Å criteria capture substantially different residue sets, producing correspondingly different pocket geometries and volume measurements. More broadly, these observations illustrate a limitation of ligand-guided residue selection for some systems: the molecular container is defined relative to the position of the bound ligand rather than the protein architecture that forms the cavity. For elongated, asymmetric, or conformationally heterogeneous pockets such as the HBV CAM site, a simple radial cutoff may unevenly sample the cavity, underrepresenting some structural features. This limitation becomes even more apparent for binding pockets capable of accommodating chemically diverse ligands. The HBV CAM site is structurally adaptable and binds compounds with a variety of sizes, shapes, and binding modes [39–41, 46, 47, 51–53]. To illustrate this behavior, the same 10 Å ligand-guided selection was applied to two independent experimental complexes: one containing AT130 in the B-site [40] and another containing DBT1 in the B-site [46] (**Figure 3E**). These ligands are distinct CAM chemotypes with different mechanisms of action. AT130 is an E-type CAM that promotes formation of stable but empty capsids, whereas DBT1 is an A-type CAM that both misdirects assembly toward aberrant products and destabilizes preformed capsids [46, 54–56]. Although the same distance criterion was applied to both ligands, the resulting residue sets differ substantially because DBT1 occupies the CAM site in a different binding mode [46] (**Figure 3E**). Thus, a ligand-guided definition makes the molecular container dependent not only on the choice of distance cutoff, but also on ligand identity and how completely the ligand fills the surrounding cavity. A Tcl script demonstrating application of the ligand-guided pocket definition approach to the HBV CAM site is provided in **Code Snippet S2**.

### Defining the binding pocket: Iterative pocket-guided approach

A pocket-guided approach defines the molecular container according to the structural elements that consistently define the cavity rather than the position of a ligand within it. This ligand-independent framework was successfully employed in a recent application of *measure volinterior* to compare CAM conformations observed in experimental structures of HBV capsids with those in flat lattices of assembly-incompetent HBV Cp [29]. Because the molecular container is defined by the protein rather than a bound ligand, this approach can be applied to both apo and ligand-bound complexes, including structures in which the ligand is incompletely resolved. For example, the HAP1-bound HBV capsid showed evidence of CAM occupancy in both the B- and C-sites [39], yet ligand density in the B-site was insufficient to resolve atomic coordinates, precluding application of a ligand-guided definition.

To localize the molecular container to the HBV CAM site, an iterative refinement procedure was employed. The objective was not simply to identify a residue set suitable for a single experimental structure, but rather to develop a molecular container that remained robust across the conformational heterogeneity of the CAM binding pocket. The four iterations shown in **Figure 4** illustrate the principal structural considerations that guided refinement of the final atom-selection (**Table 1**), progressing from an initial architecture-based definition to a molecular container that faithfully represents the pocket recognized by CAMs.

**Figure 4:**
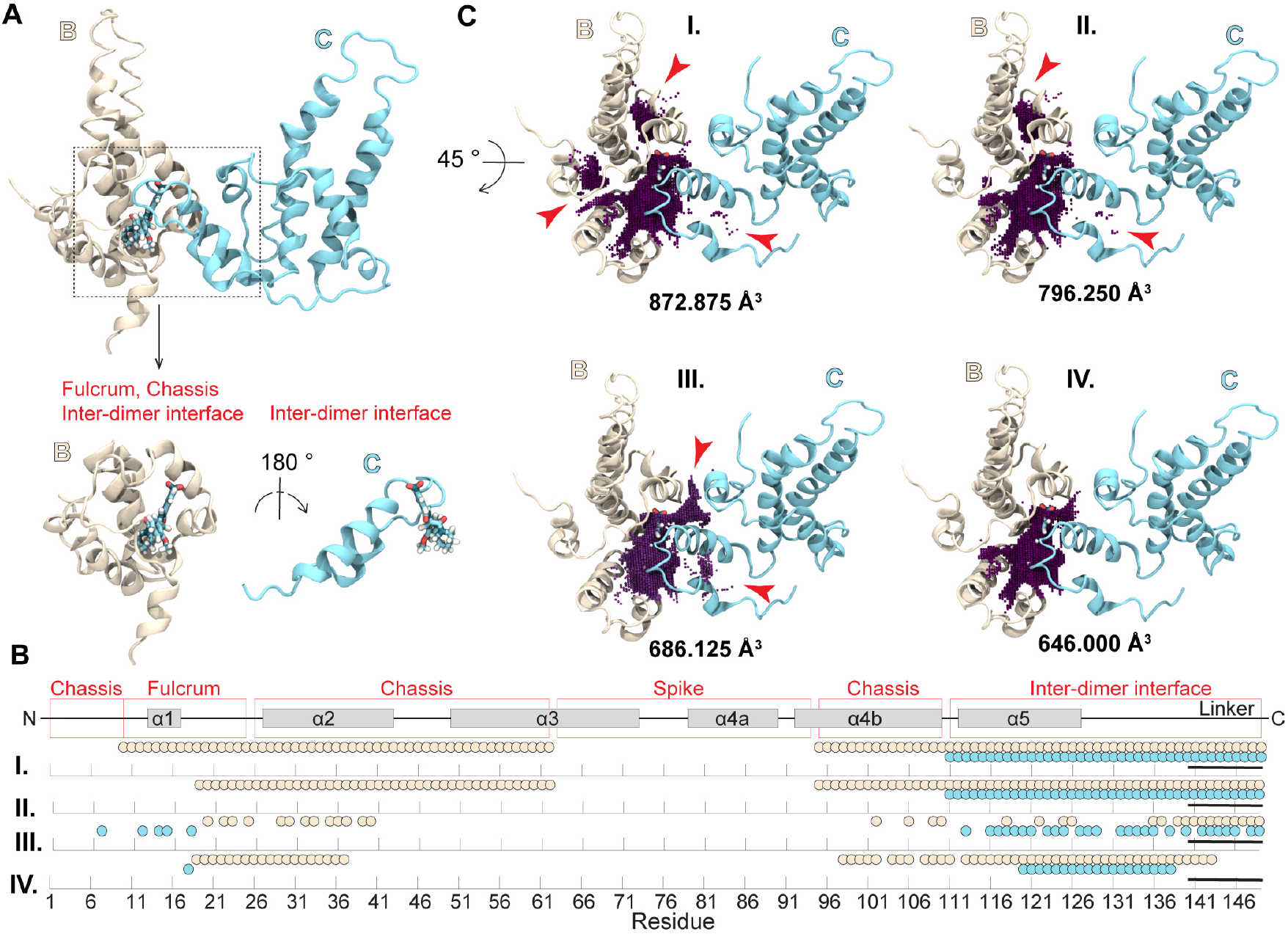
Pocket-guided definition of the CAM binding site with iterative refinement. (**A**) Structural elements of Cp149 contributing to the architecture of the CAM site, including the fulcrum, chassis, and helix–loop motif of the base chain together with the helix–loop motif of the capping chain. The helix–loop motifs from the two neighboring homodimers pack together to form the interdimer interface containing the CAM binding pocket. (**B**) Residue sets defining the molecular container across iterations of the pocket guided approach applied to the AT130-bound HBV capsid B-site [40]: (*I*) initial subdomain-based selection including residues from the structural elements contributing to the CAM site; (*II*) refinement after removal of residues producing an extraneous cavity component; (*III*) refinement guided by consensus relative SASA calculated across the quasi-equivalent CAM sites of the analyzed crystal structures; (*IV*) final refinement after adjustment of residues whose side-chain orientations generated non-pocket cavity components or unnecessarily broadened the molecular container. Modeled linker residues (residues 140–149), which lack resolved coordinates in at least one analyzed crystal structure, are indicated by a solid black line. (**C**) *measure volinterior* results obtained by applying the corresponding molecular container definitions from panel B. Each subpanel reports the calculated pocket volume together with the detected pocket geometry visualized as a three-dimensional point cloud. Red arrows indicate extraneous cavity components eliminated during successive refinement steps, illustrating progressive localization of the molecular container to the CAM site while preserving the geometry of the pocket interior. Volume calculations were performed using the optimized *measure volinterior* parameters summarized in Table 1.

#### Iteration I: Begin with a local subdomain-based definition

A logical starting point for a pocket-guided molecular container is the protein architecture surrounding the binding site. The HBV CAM site is formed by portions of the fulcrum and chassis subdomains from the Cp base chain together with the helix–loop motifs of the base and capping chains that form the interdimer interface (**Figure 4A**). Selecting residues from these structural elements provides an initial molecular container based on the organization of the protein. Application of *measure volinterior* using this initial atom-selection (**Figure 4B,I**) demonstrates that the resulting container remains overly broad. The detected pocket contains multiple enclosed regions unrelated to the CAM site, including small intrachain cavities between helix 1 and helix 4b, between helix 2 and helices 3/4b, and isolated voxels adjacent to helix 5 and the capping-chain linker (**Figure 4C,I**). These observations indicate that simply selecting structural subdomains known to contribute to the binding site can be insufficient for defining a localized molecular container that faithfully represents the pocket.

#### Iteration II: Prune residues that produce extraneous cavity components

After constructing an initial molecular container, the first refinement step is to evaluate whether the detected cavity remains localized to the binding pocket of interest. This assessment is performed by visual inspection of the cavity geometry produced by *measure volinterior*. Enclosed regions clearly unrelated to the binding site indicate that the molecular container includes residues whose molecular surface unnecessarily encloses space outside the pocket. These residues can then be removed from the atom-selection, and the cavity recalculated. For the HBV CAM site, the initial subdomain-based container produces several extraneous cavity components (**Figure 4C,I**). As an illustrative example, residues from the fulcrum contributing to the intrachain cavity between helix 1 and helix 4b are removed from the molecular container (**Figure 4B,II**). Recalculation eliminates this unwanted volume component while preserving the topology of the CAM binding pocket (**Figure 4C,II**). Although additional extraneous regions remain and require further refinement, this example illustrates the general principle of using visual inspection of the detected cavity to identify residues that unnecessarily broaden the molecular container.

#### Iteration III: Identify interface-forming residues using relative SASA

For binding pockets within protein–protein interfaces, residues included in the molecular container definition may participate in formation of the interface itself. Residues whose side chains remain solvent exposed following interface formation are less likely to contribute to the molecular surface enclosing the binding pocket than residues that become buried upon interface formation. Relative solvent-accessible surface area (SASA) [57] provides an objective means of identifying such interface-forming residues. Whenever possible, relative SASA should be evaluated across an ensemble of conformations rather than a single structure because protein–protein interfaces may be dynamic.

For the HBV CAM site, MD simulations have shown that individual interfacial contacts can vary substantially in their persistence across quasi-equivalent interfaces and ligand-bound states [34, 38]. One illustrative example is Thr109, which forms an interdimer contact in some crystal structures (e.g., PDB 1QGT [58]) but not others (e.g., PDB 2G33 [39]), while MD simulations show that this contact is populated anywhere from 0% to 57% of the time depending on the quasi-equivalent interface and CAM occupancy [34, 38]. Residues that appear buried in one conformation may therefore become solvent exposed in another. In the absence of an MD trajectory, a practical alternative is to evaluate relative SASA across an ensemble of experimentally determined structures representing different conformational states. Here, the four HBV crystal structures shown in **Figure 2**, together containing sixteen unique CAM site conformations, are used to demonstrate the approach.

Relative SASA is dimensionless and quantifies the extent to which a residue becomes buried upon formation of the interdimer interface. A Tcl script to calculate relative SASA for the HBV Cp interdimer interface is provided in **Code Snippet S3**. Importantly, SASA calculations are performed using complete Cp homodimers rather than isolated chains so that the native intradimer interface remains intact. Calculating SASA for isolated chains would incorrectly classify residues buried within the intradimer interface as solvent exposed. For each residue, side-chain SASA is first calculated for the complete homodimer containing the chain of interest, yielding *SASA*_dimer_. SASA is recalculated after adding the adjacent homodimer to form the complete dimer-of-dimers surrounding the CAM site, yielding *SASA*_dimer-of-dimers_. Relative SASA is then calculated as

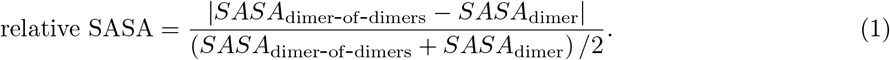

This calculation must be performed separately for both the Cp base and capping chains, comparing each homodimer in isolation (*SASA*_dimer_) with the corresponding dimer-of-dimers (*SASA*_dimer-of-dimers_, **Figure S1A**). Residues exhibiting relative SASA values greater than 0.25 are classified as interface-forming residues [57] (**Figure S1B**).

A consensus residue set arises from pooling all residues exceeding the relative SASA threshold in at least one of the quasi-equivalent CAM site conformations in the analyzed crystal structures. (**Figure 4B,III**). Compared with the previous iteration, this refinement of the molecular container removes several residues that make little contribution to the protein–protein interface while identifying additional residues involved in interface formation. However, interface formation alone does not uniquely define the CAM site. Some residues contribute to the interdimer interface without contributing to the molecular surface enclosing the binding pocket, leading to extraneous cavity components (**Figure 4C,III**). Consequently, although relative SASA provides an objective and highly effective refinement criterion, additional structural refinement is still required to produce a molecular container definition specifically localized to the CAM site.

#### Iteration IV: Refine the molecular container using side-chain orientation

Finally, refinement should be guided by visual inspection of the detected cavity together with the orientations of individual residues defining the molecular surface. Although the relative SASA analysis successfully identifies residues participating in the interdimer interface, not every interfacial residue contributes equally to the architecture of the CAM site. Side chains directed away from the site may unnecessarily enlarge the molecular container or generate isolated cavity components outside the pocket. Residues Lys96 and Phe97 are excluded because their side chains project away from the binding pocket, whereas Arg98 is retained because it projects toward the fulcrum and closes intrachain gaps that define the pocket cavity (**Figure S2A**). Additional helix 4b residues, including Phe103, Ala107, and Arg112, are removed because their side chains contribute isolated voxels outside the pocket (**Figure S2B**). In the capping chain, only Phe18 is retained from the additional fulcrum residues identified by the relative SASA analysis because its side chain projects toward the pocket interior and improves closure of the detected cavity without introducing extraneous enclosed regions (**Figure S2C**). Residues 111–120 are excluded because they promote isolated voxels adjacent to helix 5 and the linker. These refinements produce the final molecular container definition (**Figure 4B,IV**), yielding the cleanest and most localized representation of the CAM binding pocket (**Figure 4C,IV**). This atom-selection is used throughout the remainder of the study as part of the optimized parameter set recommended for the HBV CAM site (**Table 1**).

Although development of a robust pocket-guided molecular container requires substantially greater effort than applying a simple ligand-distance cutoff, the resulting definition is independent of ligand identity, binding mode, and occupancy state. Consequently, the same atom-selection can be applied consistently to apo structures, ligand-bound complexes containing different CAM chemotypes, and conformational ensembles generated by MD simulations without redefining the binding pocket for each new structure. For highly dynamic systems such as the HBV CAM site, this consistency is particularly valuable because it ensures that changes in measured pocket geometry reflect genuine conformational fluctuations rather than differences in the definition of the molecular container.

The four iterations presented here illustrate one possible refinement pathway rather than a universally prescribed procedure. During development of the HBV CAM site protocol, more than sixteen intermediate residue sets were evaluated before arriving at the final molecular container. Each refinement was motivated by objective structural information, including protein architecture, interface burial, and side-chain orientation, and subsequently evaluated by visual inspection of the detected cavity geometry. Although considerably fewer iterations may be sufficient when analyzing a single experimental structure, optimization against conformational ensembles benefits from development of a consensus molecular container capable of faithfully representing the full range of pocket heterogeneity. More generally, molecular container development should be viewed as an iterative, data-driven process in which refinement of the atom-selection and evaluation of the resulting cavity geometry proceed hand-in-hand.

### Defining the binding pocket: Impact of disordered and unresolved residues

Before finalizing a molecular container, it is important to evaluate whether residues contributing to the binding pocket are intrinsically disordered or incompletely resolved in the available structural data. Such regions present a unique challenge because their conformations may be highly variable across structural ensembles or absent altogether from experimental structures. Consequently, inclusion of these residues can introduce uncertainty into the detected cavity geometry, particularly when comparing structures with different chain lengths. The HBV CAM site provides an instructive example of this problem.

The B- and C-sites extracted from the AT130-bound capsid represent challenging test systems, owing to the relatively low resolution of the structure (4.2 Å) and incomplete definition of the Cp149 linker region [40]. The linker (residues 140–149) connecting helix 5 to the C-terminal domain (residues 150-183 in native Cp) is highly flexible and is only partially resolved by experiments. Consequently, different experimental structures contain different extents of the linker, requiring computational structural prediction to rebuild complete Cp149 [34, 42, 43]. To evaluate the influence of linker definition on the molecular container, *measure volinterior* calculations were performed using residue selections containing either the experimentally resolved linker or the reconstructed full-length linker (**Figure 5**). The resulting cavity geometries demonstrate that inclusion of alternative linker lengths or conformations can substantially alter the enclosure of the CAM pocket and consequently the calculated volume. Thus, even when the remainder of the molecular container is unchanged, uncertainty associated with a flexible structural element can become a dominant source of variability in cavity characterization.

**Figure 5:**
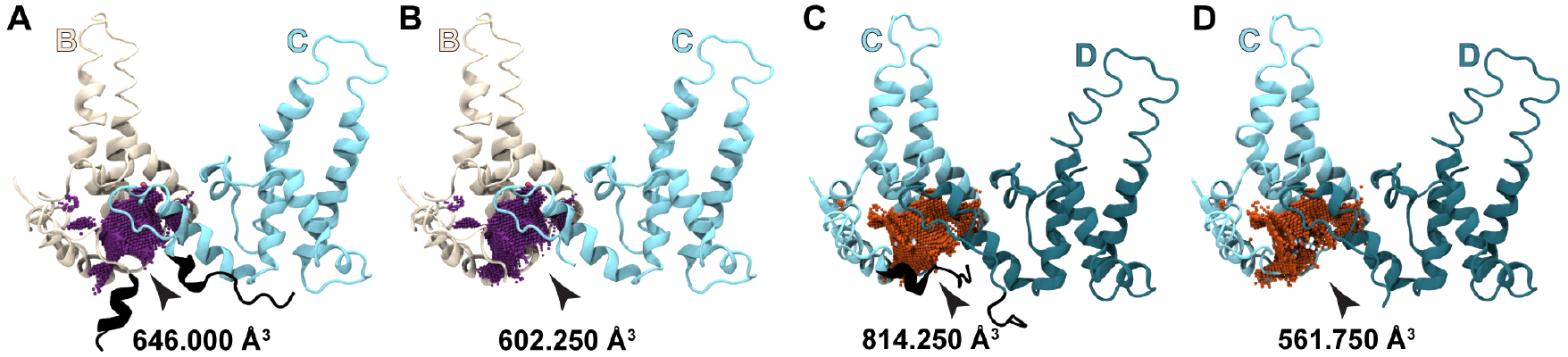
Linker residues influence CAM binding site geometry and volume. Comparison of CAM pocket geometries and calculated volumes obtained using *measure volinterior* for the AT130-bound quasi-equivalent B-site with (**A**) reconstructed linker residues (full-length Cp149 model) and (**B**) only the experimentally resolved linker residues present in the original crystal structure. The corresponding comparison for the quasi-equivalent C-site is shown in (**C**) and (**D**), respectively. Residues modeled to reconstruct the unresolved linker are shown in black. Although the molecular container is otherwise identical, inclusion of the modeled linker substantially alters the detected pocket geometry and calculated volume, illustrating the sensitivity of cavity characterization to unresolved and highly flexible structural elements. Volume calculations were performed using the optimized *measure volinterior* parameters summarized in Table 1.

This issue is particularly important for dynamic structural ensembles. Previous all-atom MD simulations of the intact Cp149 capsid showed that residues beyond Thr142 exhibit C*α* root-mean-square fluctuations exceeding 4 Å, indicating substantial conformational heterogeneity in the linker region [42]. As a result, including the full-length linker in the molecular container will cause the detected pocket geometry to fluctuate primarily in response to linker motion rather than changes in the binding pocket itself. For the HBV CAM site, these observations motivated truncation of the linker at residue 143 in the Cp base chain and residue 138 in the capping chain, yielding the final molecular container shown in **Figure 4B,IV**. More generally, when highly flexible or incompletely resolved structural elements contribute to a binding pocket, it may be advantageous to exclude the most mobile portions of those regions while retaining only residues that make consistent contributions to the molecular surface enclosing the cavity. The appropriate truncation point will depend on the structural system and should be guided by the extent to which the flexible region influences the detected pocket geometry across the conformational ensemble.

### Constructing the molecular surface: Choosing QuickSurf parameters

The second step in applying *measure volinterior* to a binding pocket is construction of the molecular surface using the QuickSurf representation. For the HBV CAM site, the goal is to select the Radius Scale, Isovalue, and Grid Spacing parameters such that the localized molecular container faithfully represents the structural features surrounding the binding pocket while minimizing artifacts that distort the detected cavity geometry. The molecular surface generated by QuickSurf can be viewed as a mold that defines the envelope of the protein, whereas the cavity detected by *measure volinterior* corresponds to the cast of the enclosed negative space produced by that mold. Consequently, QuickSurf parameterization should be guided by the geometry, topology, and continuity of the detected cavity rather than by visual appearance of the molecular surface of the protein alone. Because Radius Scale, Isovalue, and Grid Spacing collectively determine the molecular surface, these parameters must be optimized together rather than independently. Furthermore, because the molecular surface is constructed from the residues comprising the molecular container, refinement of the QuickSurf parameters may necessitate reevaluation of the atom-selection used to define the pocket.

The QuickSurf representation is fundamentally constructed from atomic coordinates and their associated atomic radii. As discussed in the Methods, the protocol described here uses the default VMD vdW radii. Consequently, the molecular surface generated by QuickSurf represents an expansion upon the vdW sphere representation (VMD’s VDW drawing method) and remains inherently dependent on the values of the atomic radii used during surface construction. Holding Radius Scale, Isovalue, and Grid Spacing constant, modification of the underlying atomic radii alters the resulting molecular surface and therefore changes the volume and topology of the detected cavity (**Figure 6A**). Furthermore, as discussed in Methods, reproducibility of this protocol requires use of the GPU implementation of *measure volinterior*, as the CPU and GPU versions of the code produce different results even when identical QuickSurf parameters are used (**Figure 6A–B**).

**Figure 6:**
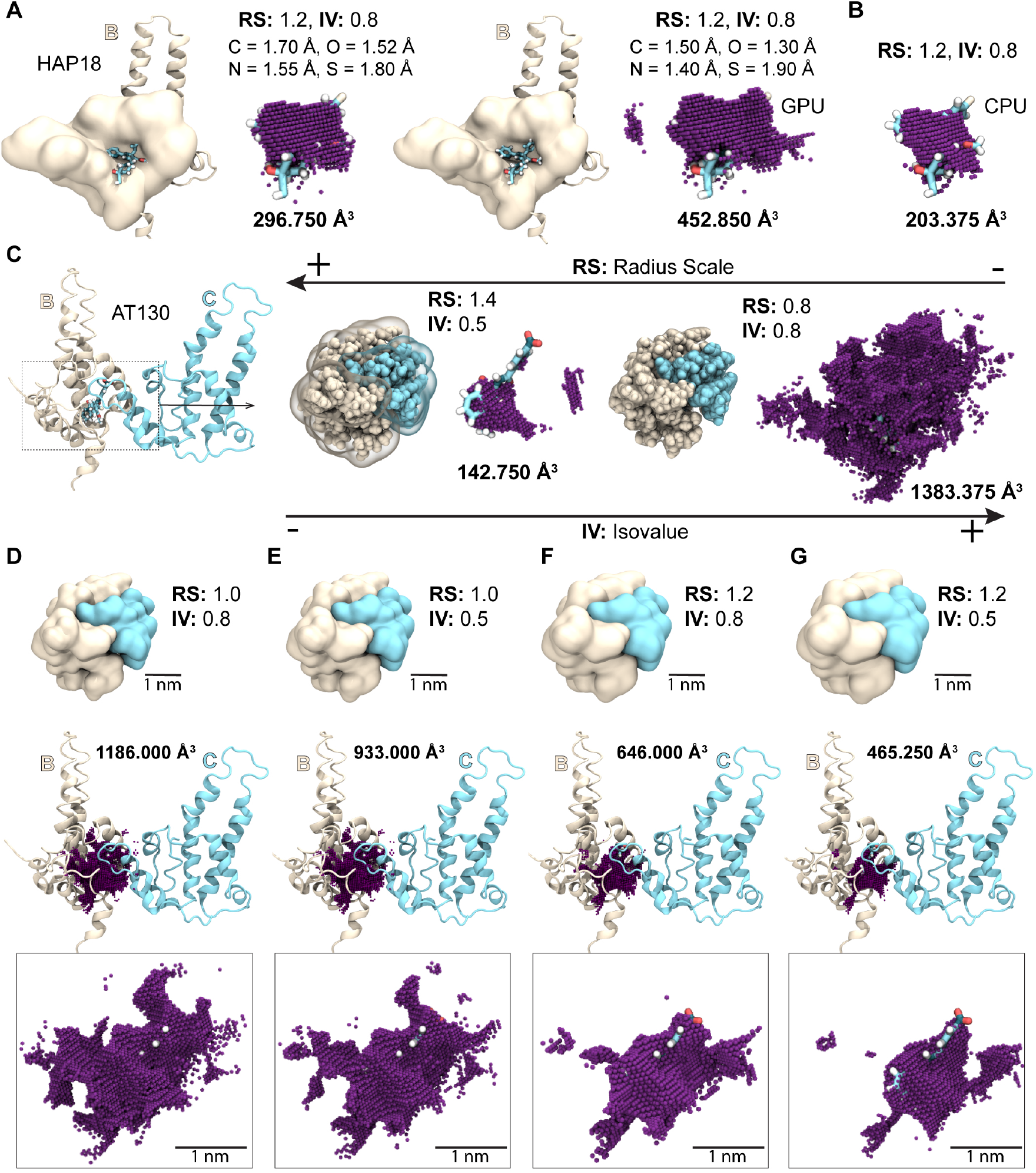
Optimizing the Radius Scale and Isovalue parameters. (**A**) Choice of atomic radii set influences QuickSurf molecular surface and resulting CAM pocket geometry detected by *measure volinterior*. (**B**) Given equivalent parameters, results obtained using the CPU versus GPU implementations (as in panel A, right) are not equivalent. (**C**) Effect of extreme Radius Scale (RS) and Isovalue (IV) combinations on the molecular surface and detected CAM pocket geometry. Excessively bulky surfaces can intrude into the binding pocket, whereas excessively thin surfaces may underrepresent the protein itself. (**D–G**) Pocket geometries obtained using Radius Scale and Isovalue combinations within the recommended calibration range. Increasing Radius Scale improves localization of the detected cavity; the combination of RS = 1.2 and IV = 0.8 provides the cleanest representation of the HBV CAM pocket. Unless otherwise noted, volume calculations were performed using the optimized *measure volinterior* parameters summarized in Table 1.

Although Radius Scale, Isovalue, and Grid Spacing are interdependent, Radius Scale provides the most direct control over the effective bulkiness of the molecular surface and therefore serves as a useful starting point for optimization. **Figure 6C** demonstrates that Radius Scale exerts a substantially larger influence on surface geometry than Isovalue. At large Radius Scale values, such as 1.4 Å, the surface becomes excessively bulky and intrudes into the CAM pocket, artificially reducing the detected cavity volume. In contrast, small Radius Scale values, such as 0.8 Å, produce surfaces that contract toward the atomic centers and may even fall within the envelope defined by the vdW spheres themselves. Under these conditions, portions of the protein volume are incorrectly classified as pocket interior, resulting in artificially enlarged and rugged cavity geometries.

For the HBV CAM site, Radius Scale values between 1.0–1.2 Å provided a practical range for calibration. Within this range, the QuickSurf surface extends only modestly beyond the underlying vdW envelope of the protein, providing a reasonable approximation of the steric boundary experienced by a ligand. Such expansions are small relative to the characteristic separation of noncovalently interacting atoms, which typically approach one another no closer than approximately 3–4 Å. Consequently, a molecular surface extending a few tenths of an Ångstrom beyond the vdW envelope is unlikely to substantially distort the physically relevant pocket geometry while helping to eliminate small surface discontinuities that can complicate cavity detection. Values above this range should be used cautiously because the molecular surface begins to occupy space that would otherwise belong to the pocket interior, whereas values below this range tend to underestimate the physical bounds of the protein surface.

After establishing a suitable Radius Scale range, Isovalue can be used to fine-tune the molecular surface. The effect of varying Isovalue within the recommended Radius Scale range is shown in **Figure 6D–G**. At a Radius Scale of 1.0 Å, the detected CAM pocket remains fragmented and contains numerous isolated voxels regardless of the tested Isovalue (**Figure 6D–E**). Increasing the Radius Scale to 1.2 Å improves delineation of the pocket and reduces extraneous cavity components, permitting more meaningful evaluation of Isovalue (**Figure 6F–G**). Under these conditions, an Isovalue of 0.5 produces a cavity that is artificially small, whereas an Isovalue of 0.8 yields a cleaner and more faithful representation of the CAM site. Based on these observations, a Radius Scale of 1.2 Å and an Isovalue of 0.8 were selected as optimal parameters.

In contrast to Radius Scale and Isovalue, Grid Spacing primarily controls the spatial resolution of both the molecular surface and the resulting cavity representation. Smaller Grid Spacing values capture finer structural detail but increase computational cost, whereas larger values reduce resolution and can obscure important features of the pocket geometry. For large molecular containers such as intact capsids or vesicles, Grid Spacing values exceeding 1.0 Å may preserve sufficient information to characterize the massive enclosed volume. For small protein cavities, however, finer spatial sampling is generally required to resolve atomistic features of the pocket. **Figure 7** illustrates the effects of coarse and fine Grid Spacing values on both isosurface and point-cloud representations of the HBV CAM pocket. As Grid Spacing increases, individual voxels become larger, leading to progressive loss of geometric detail. Grid Spacing values between 0.5–1.0 Å generally provide a practical balance between resolution and computational efficiency for binding pockets. For the HBV CAM site, a Grid Spacing of 0.5 Å was chosen to maximize fidelity of the detected cavity geometry. At this resolution, each voxel corresponds to 0.125 Å^3^, explaining why all pocket volumes reported herein are integer multiples of this fundamental unit of three-dimensional space.

**Figure 7:**
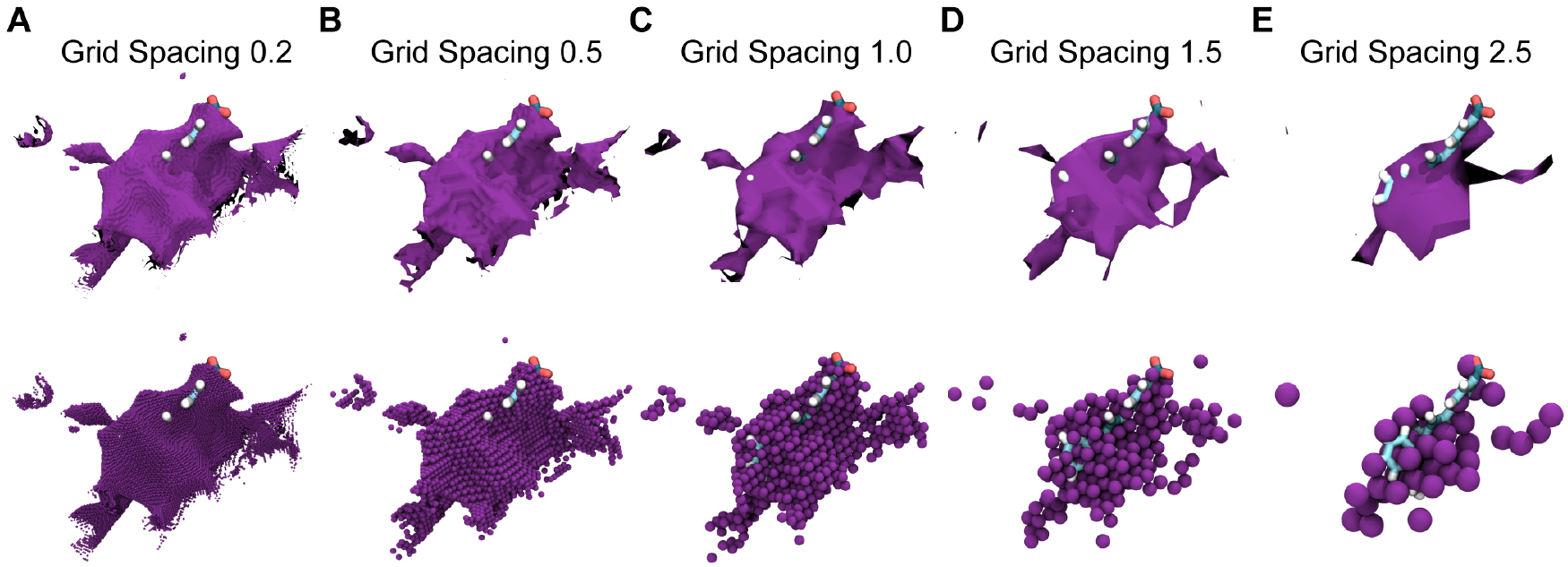
Optimizing the Grid Spacing parameter. CAM pocket geometries produced using Grid Spacing values of (**A**) 0.2 Å, (**B**) 0.5 Å, (**C**) 1.0 Å, (**D**) 1.5 Å, and (**E**) 2.5 Å. For each Grid Spacing, the upper panel shows the pocket geometry visualized as an isosurface (with Isovalue 0.8) using the volumetric map output by *measure volinterior*, while the lower panel shows the corresponding three-dimensional point-cloud representation reconstructed from the same volumetric data. Decreasing Grid Spacing preserves finer structural detail at increased computational cost, increasing larger Grid Spacing produces progressively coarser pocket representations. The close correspondence between the isosurface and point-cloud visualizations demonstrates that the reconstructed point cloud faithfully reproduces the topology of the underlying volumetric map. Unless otherwise noted, calculations were performed using the optimized *measure volinterior* parameters summarized in Table 1.

Importantly, multiple combinations of Radius Scale, Isovalue, and Grid Spacing may generate molecular surfaces that produce acceptable cavity representations for a given system. Consequently, a unique optimal parameter set need not exist. Nevertheless, parameter selection should be guided by the resulting cavity geometry rather than by visual inspection of the molecular surface alone. Because the molecular surface depends on both the QuickSurf parameters and the underlying atom-selection defining the binding pocket, optimization of these interrelated components should be viewed as an iterative process in which refinement of one may necessitate reevaluation of the other.

Although the Number of Rays parameter does not influence construction of the molecular surface itself, it determines how thoroughly that surface is sampled during ray-casting calculations and therefore affects characterization of the detected cavity. Number of Rays controls the angular sampling used to evaluate local enclosure around each voxel and thus influences the accuracy with which pocket boundaries are detected. The original *measure volinterior* publication recommended values of *N*_rays_ ≥ 32 [19], and for large molecular containers like intact virus capsids, this value often provides a reasonable compromise between accuracy and computational performance. In contrast, the HBV CAM site contains narrow features, irregular local geometry, and partially occluded regions that benefit from denser angular sampling. Based on the present analyses, *N*_rays_ = 64 provides a suitable balance between computational efficiency and robust characterization of pocket geometry, particularly when applied over a large conformational ensemble. In general, once the surrounding molecular surface becomes sufficiently sampled, the resulting cavity topology and volume calculations become largely invariant to further increases in Number of Rays. Consequently, values of *N*_rays_ ≫ 64 are unlikely to provide meaningful improvements for characterization of most binding pockets while unnecessarily incurring additional computational cost.

### Defining the pocket boundary: Choosing the occlusion threshold

The third step in applying *measure volinterior* to a binding pocket is selection of the occlusion threshold defining the effective boundary of the cavity across its solvent-accessible opening. For the HBV CAM site, the goal is to choose an occlusion threshold that preserves the physically relevant topology of the binding pocket while excluding weakly occluded regions that do not contribute meaningfully to the cavity interior. Because the CAM site is not a fully enclosed container, characterization of its volume requires use of fuzzy-boundary detection [19]. Under this approach, each voxel is assigned an occlusion score reflecting the fraction of rays that intersect the surrounding molecular surface, and an occlusion threshold is subsequently applied to determine whether the voxel is retained as part of the detected cavity. Consequently, the selected threshold exerts direct control over the geometry and topology of the resulting pocket representation.

**Figure 8** illustrates the effect of the occlusion threshold on the detected CAM site when the molecular container (pocket-defining atom-selection) and molecular surface (QuickSurf parameters) are fixed at their optimized values (**Table 1**). At lower occlusion thresholds of 0.7 and 0.8, the detected pocket incorporates solvent-exposed regions beyond the core cavity (**Figure 8A-B**). Although these voxels satisfy the selected occlusion criterion, they contribute little to the physically relevant volume available for ligand binding and artificially inflate the reported pocket size. At an occlusion threshold of 0.9, the detected cavity is mostly continuous and localized to the ligand binding pocket (**Figure 8C**). Although some some extraneous voxels remain, the volume of space accessible to the ligand is reasonably represented and controlled primarily by the orientations of pocket-defining side-chains. A more restrictive threshold, such as 0.99, might be expected to provide a cleaner definition of the cavity by retaining only voxels with a very high probability of belonging to the pocket interior (**Figure 8D**). Instead, increasing the threshold disrupts the detected topology, fragmenting the cavity and artificially excluding portions of the pocket clearly accessible to the ligand. Thus, by biasing the measurement toward only the most deeply buried regions of the binding site, excessively restrictive occlusion thresholds can underrepresent the pocket and yield cavity geometries that no longer reasonably reflect the space accessible to a bound ligand.

**Figure 8:**
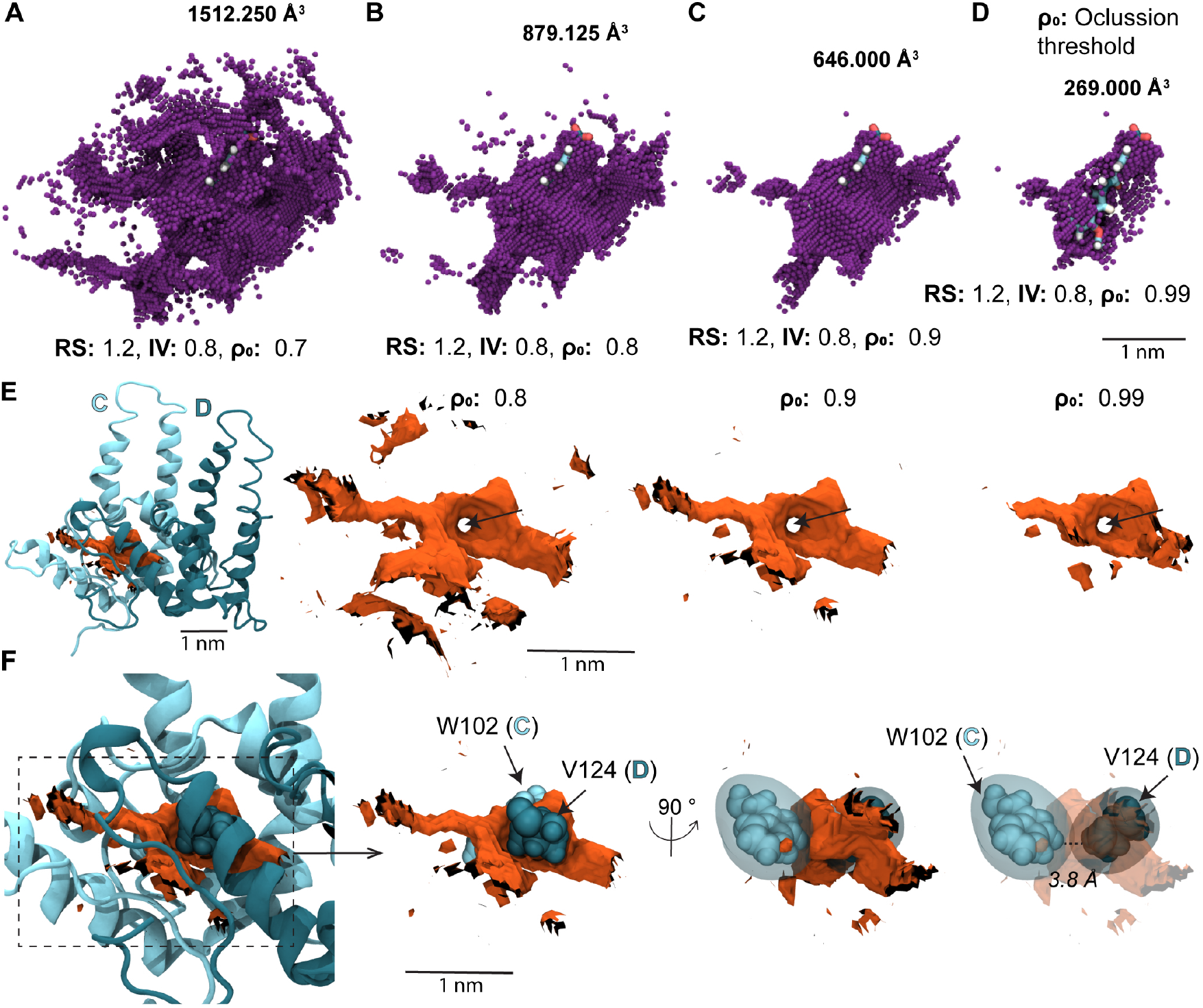
Optimizing the occlusion threshold. (**A–D**) CAM site topologies and corresponding volumes detected by *measure volinterior* for the AT130-bound B-site using occlusion thresholds of (**A**) 0.7, (**B**) 0.8, (**C**) 0.9, and (**D**) 0.99. Lower thresholds retain weakly occluded solvent-accessible regions, whereas increasingly restrictive thresholds reduce the detected volume and eventually fragment the pocket topology. (**E**) Example of an acceptable local discontinuity in the detected cavity topology for the apo-form C-site. Pocket topologies are shown using occlusion thresholds of *ρ*_0_ = 0.8, 0.9, and 0.99. The persistent local discontinuity is indicated by the black arrow. (**F**) Close-up view showing that the discontinuity arises from the underlying protein geometry rather than the choice of occlusion threshold. The molecular surfaces surrounding Trp102 of chain C and Val124 of chain D merge to occlude the intervening space, producing a small excluded region (“donut hole”) within the detected pocket. Such features should be retained when they reflect the physical organization of the molecular surface rather than artifacts of parameter selection. Unless otherwise noted, volume calculations were performed using the optimized *measure volinterior* parameters summarized in Table 1.

Importantly, not all discontinuities in the detected cavity topology are artifacts that should be eliminated through adjustment of the occlusion threshold. While isolated voxels and disconnected cavity fragments unrelated to ligand binding are generally undesirable, local discontinuities may also arise from the physical organization of the protein. Local side-chain packing can partition the cavity and create void spaces that are inaccessible to the ligand. Such features may fluctuate across a structural ensemble as side-chain conformations change, reflecting genuine conformational dependence of the binding-site geometry. An example is shown in **Figure 8E-F**, where a local discontinuity in the detected CAM pocket persists across a range of occlusion thresholds. Inspection of the underlying protein structure reveals that this feature arises from two interacting side-chains whose combined molecular surface occludes the intervening space. Although the corresponding vdW spheres appear sufficiently separated to suggest the presence of a continuous cavity, application of the optimized QuickSurf parameters causes the surfaces surrounding these residues to merge, producing a small excluded region (“donut hole”) within the detected pocket topology. This feature therefore represents a physically meaningful consequence of the selected molecular container, molecular surface, and local protein conformation rather than an artifact of the occlusion threshold.

For the HBV CAM site, an occlusion threshold of 0.9 provides the best balance between preserving a continuous pocket topology and excluding weakly occluded solvent-accessible regions. More generally, these results suggest that occlusion thresholds between 0.7–0.9 provide a practical starting range for evaluation of other dynamic binding pockets. The optimal value for a system will depend on the geometry of the cavity and should ultimately be guided by the resulting pocket topology rather than by the numerical value of the threshold itself. Importantly, the occlusion threshold cannot be optimized independently of the molecular container and molecular surface. Because occlusion scores are calculated by evaluating ray intersections with the surface generated for the container residues, refinement of the occlusion threshold may alter the apparent suitability of the underlying atom-selection and QuickSurf parameters. Consequently, optimization of the pocket definition should be viewed as an iterative process in which refinement of any one of these three coupled parameterization components may necessitate reevaluation of the others.

### Final considerations for reproducible pocket volume comparisons

Beyond consistent parameters, reproducible application of *measure volinterior* requires that structures be analyzed in a common spatial orientation. This consideration arises because *measure volinterior*, like many pocket-volume algorithms, represents the molecular surface on a three-dimensional Cartesian voxel grid [19]. Although the detected cavity is ultimately determined by the molecular surface, classification of individual voxels depends on how that surface intersects the grid. Consequently, rigid-body rotation of an otherwise identical structure can produce small differences in the detected cavity geometry and calculated volume. In contrast, rigid-body translation does not introduce the same effect. Because the voxel grid is generated by fitting an orthorhombic bounding box around the selected molecular container, translating the structure simply translates the fitted grid together with it. The relative placement of atoms within the grid therefore remains unchanged, preserving the resulting voxel classification.

To evaluate the magnitude of orientation-dependent systematic error, the quasi-equivalent B-site CAM pocket from the AT130-bound capsid [40] was subjected to rigid-body rotations about the Cartesian *x*^, *y*^, and *z*^ axes in 5*^◦^* increments over a full 360*^◦^* (**Figure 9A**). Pocket volumes were recalculated for each orientation using the optimized parameters summarized in **Table 1**. Small but reproducible differences in the calculated pocket volume were observed for rotations about all three axes (**Figure 9B**). Comparison of the corresponding cavity geometries showed that the overall topology of the binding pocket remained unchanged, with differences localized primarily to voxels near the pocket opening and cavity periphery where small changes in voxel assignment occur (**Figure 9C**). For the AT130 B-site, the maximum orientation-dependent difference was approximately 49 Å^3^. A similar analysis across all quasi-equivalent CAM sites shown in **Figure 2** demonstrates that the maximum orientation-dependent variation remains below approximately 70 Å^3^ for every structure examined (**Figure 9D**). These differences are systematic rather than stochastic, and although modest relative to the overall pocket volume, can still complicate quantitative comparison of closely related conformations.

**Figure 9:**
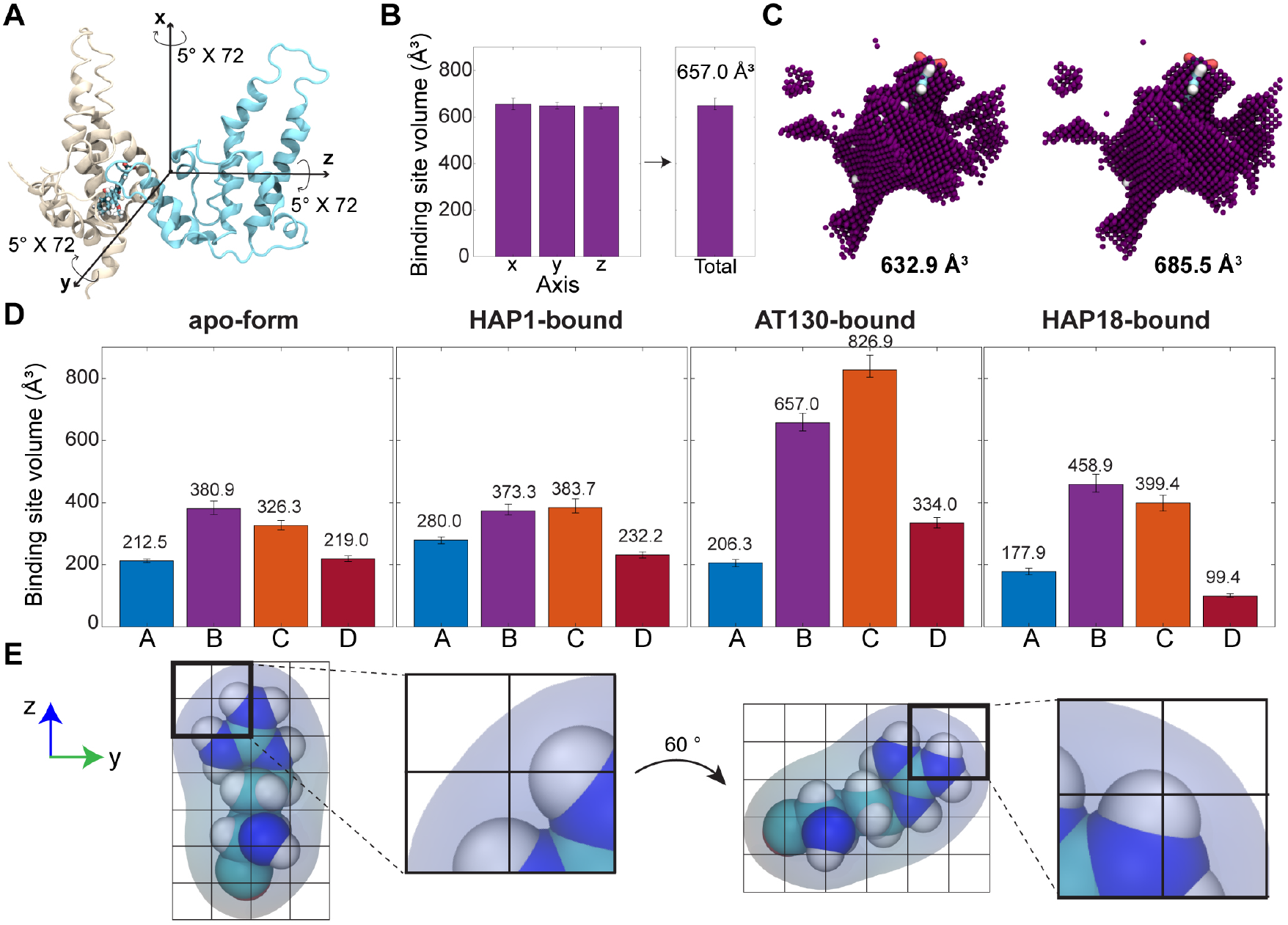
Rigid-body rotation introduces systematic variation in pocket volume calculated by *measure volinterior*. (**A**) The quasi-equivalent B-site CAM pocket from the AT130-bound capsid was subjected to independent rigid-body rotations about the Cartesian *x*^, *y*^, and *z*^ axes. For each axis, 72 successive rotations of 5*^◦^* were applied to complete a full 360*^◦^* rotation, and the pocket volume was recalculated after each rotation. (**B**) Average pocket volumes obtained for each rotational ensemble. Error bars indicate the minimum and maximum volume values observed over the full rotation about each axis. The cumulative average across all three rotational ensembles is also shown (657.0 Å^3^). (**C**) Detected pocket geometries corresponding to the minimum- and maximum-volume conformations observed during the rotational analysis. Although the overall pocket geometry remains qualitatively unchanged, rigid-body rotation produces a systematic volume difference of approximately 49 Å^3^ owing to changes in voxel assignment near the molecular surface. (**D**) Orientation dependence across the complete HBV crystal benchmark set. Average pocket volumes are shown for all four quasi-equivalent CAM sites extracted from the apo, HAP1-bound, AT130-bound, and HAP18-bound capsids following the rotational analysis described in panel A. Error bars represent the minimum and maximum volume values obtained for each pocket, demonstrating that orientation-dependent variation remains below approximately 70 Å^3^ across all analyzed structures. (**E**) Schematic illustrating the origin of the orientation dependence. Rigid-body rotation changes the dimensions of the molecular container projected onto the Cartesian axes and therefore the dimensions of the fitted voxel grid. Consequently, atoms occupy different positions relative to voxel boundaries, leading to small differences in voxel classification and calculated pocket volume despite an unchanged protein conformation. Volume calculations were performed using the optimized *measure volinterior* parameters summarized in Table 1.

The origin of this behavior is illustrated schematically in **Figure 9E**. Rigid-body rotation changes the dimensions of the molecular container projected onto the Cartesian axes and therefore alters the dimensions of the fitted voxel grid. As a consequence, atoms occupy different positions relative to voxel boundaries, leading to small changes in the classification of voxels near the molecular surface even though the underlying protein conformation is unchanged. Similar orientation-dependent behavior has been reported for other grid-based pocket volume methods [59, 60]. Accordingly, structures should ideally be aligned to a common reference before applying *measure volinterior* whenever quantitative comparison of pocket volumes is desired. For the HBV CAM site, the asymmetric unit of the apo-form capsid structure [39] was translated to the origin and aligned to its principal axes using the VMD Orient package [20]; all remaining structures were aligned to this reference using the relatively rigid Cp149 chassis (residues 1–10, 26–62, and 95–110, **Figure 1a**) prior to volume calculation. Analogous alignment procedures should ideally be incorporated into workflows applying *measure volinterior* to structural ensembles in order to minimize orientation-dependent systematic error.

### Validating optimized parameters over conformational ensembles

Once a satisfactory *measure volinterior* parameter set (atom-selection, QuickSurf representative, occlusion threshold) has been established, it should be validated against a representative conformational ensemble before large-scale application to MD trajectories. The purpose of this validation step is not necessarily to re-optimize the parameters across millions of conformations, but rather to confirm that the selected parameterization continues to produce physically meaningful cavity geometries throughout the conformational landscape sampled by the binding pocket. Validation also provides an opportunity to identify systematic failures that would warrant revisiting the molecular container definition, QuickSurf parameters, or occlusion threshold before analyzing the complete ensemble. For long timescale MD simulations, this assessment can be performed efficiently using a strided subset of the trajectory before applying the validated parameters to the full dataset. The HBV CAM site provides an instructive example of this validation strategy.

Here, the optimized parameter set was evaluated using previously a published 1-*µ*s all-atom MD simulation of the HAP1-bound HBV capsid [34]. Three million conformations were analyzed for both the quasi-equivalent B- and C-sites. Although the corresponding crystal structure contains HAP1 bound in both of these pockets, atomic coordinates could only be resolved for the C-site ligand. Consequently, the MD simulation contains HAP1 only in the C-site pockets, meaning that differences between the B- and C-site ensembles may reflect both quasi-equivalence and ligand occupancy. The resulting volume distributions demonstrate that the CAM site samples a substantially broader range of pocket conformations than represented by the corresponding crystal structure (**Figure 10A**). Although the B- and C-sites exhibit similar initial pocket volumes (**Figure 2**), their distributions diverge during the simulation. These observations emphasize the importance of validating parameter performance using relaxed conformational ensembles rather than relying solely on the experimental structures.

**Figure 10:**
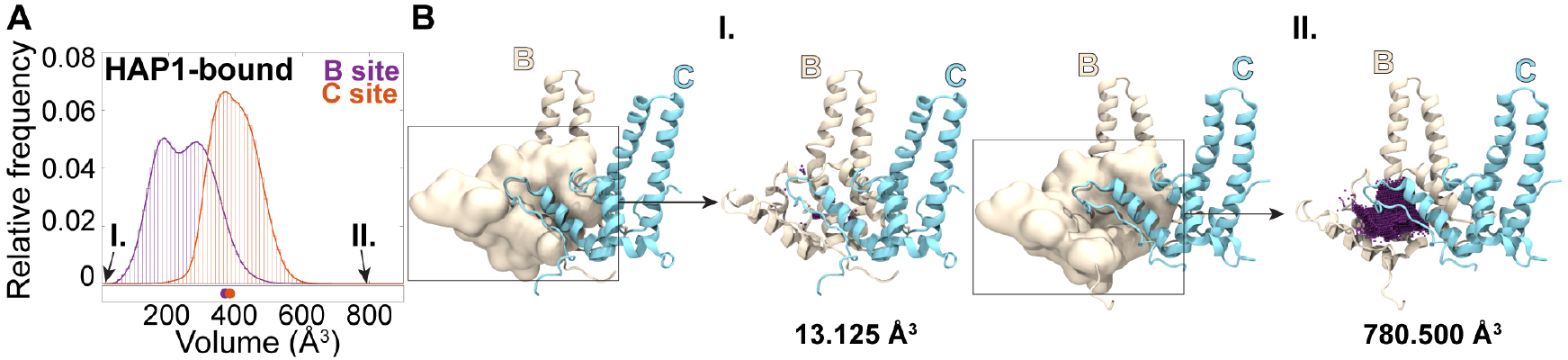
Validation of optimized *measure volinterior* parameters over an MD-derived CAM binding site ensemble. (**A**) Pocket volume distributions calculated for the quasi-equivalent B-site (purple) and C-site (orange) CAM pockets from a previously published 1-*µ*s MD simulation of the HAP1-bound HBV capsid [34]. Each ensemble comprises three million trajectory frames, for a total of six million analyzed CAM-site conformations. Colored dots beneath the distributions indicate the pocket volumes of the starting crystal structure conformations (Figure 2). The distributions illustrate that the CAM site samples a substantially broader range of pocket conformations than represented by the initial experimental structure. (**B**) Representative conformations from the tails of the HAP1-bound B-site volume distribution corresponding to the minimum (I) and maximum (II) measured pocket volumes. Although these conformations represent the extremes of the sampled conformational landscape, the detected cavity remains localized to the CAM site, illustrating the robustness of the optimized parameter set (summarized in Table 1) across highly contracted and expanded pocket states.

A practical strategy for validating optimized parameters is to inspect representative conformations from the tails of the volume distribution. These structures provide stringent test cases for determining whether unusually small or large measured volumes reflect genuine changes in pocket geometry or failures of the selected parameterization. For the HAP1-bound B-site ensemble, the minimum- and maximum-volume conformations were examined in detail (**Figure 10B**). The minimum-volume conformation corresponds to local closure of the CAM pocket, in which residues from helix 5 of the Cp capping chain project into the groove formed by the base chain. Conversely, the maximum-volume conformation represents a substantially expanded CAM pocket. Although the measured volume is nearly twice that of the corresponding crystal structure, the detected cavity remains localized to the CAM binding site and does not fragment into disconnected cavity components or extend into unrelated regions of the protein. These observations indicate that the optimized parameter set remains robust across the conformational extremes sampled by the MD simulation while continuing to faithfully describe the binding pocket.

## Conclusions and outlook

The *measure volinterior* algorithm provides a flexible approach for characterization of binding pockets, but meaningful application requires careful selection of the parameters governing the molecular container, molecular surface, and pocket boundary. Because these three components are inherently coupled, parameter optimization cannot be reduced to independent adjustment of individual numerical values. Instead, development of a robust parameterization should be viewed as an iterative process in which the resulting cavity geometry is evaluated alongside the underlying protein structure until a physically meaningful representation of the binding pocket is obtained.

The HBV CAM site provides a particularly demanding test case because the pocket is highly dynamic, partially solvent accessible, and formed at a protein–protein interface. Using this system as an illustrative example, a practical workflow was developed for parameterizing *measure volinterior*, resulting in the standardized parameter set summarized in **Table 1**. Given their demonstrated robustness and reproducibility, these optimized parameters are proposed as a community standard for characterization of the HBV CAM site, enabling direct and reproducible comparisons of pocket volume, geometry, and topology across independent experimental and computational studies.

More generally, the examples presented here suggest qualitative criteria for evaluating parameter suitability in other binding pockets. The objective of parameter optimization is not to maximize or minimize the calculated pocket volume, but rather to obtain a cavity representation that faithfully reflects the physical space enclosed by the protein. A suitable parameterization should therefore (i) remain localized to the binding pocket of interest without extending into unrelated cavities elsewhere in the protein, (ii) preserve a continuous cavity whenever the underlying protein architecture defines a continuous ligand-accessible space, (iii) closely approximate the steric boundary of the protein without either encroaching on the pocket interior or penetrating into the protein volume, and (iv) produce changes in pocket geometry that can be interpreted in terms of underlying structural rearrangements rather than artifacts introduced by parameter selection. When these criteria are satisfied across representative conformations spanning the expected conformational landscape, the resulting parameter set can be considered suitable for binding pocket characterization.

Although the optimal parameter values will ultimately depend on the structural characteristics of a particular system, the analyses presented here provide practical starting points for development of new parameterizations. For binding pockets, Radius Scale values of 1.0–1.2 Å, Isovalues of 0.5–0.8, Grid Spacing values of 0.5–1.0 Å, occlusion thresholds of 0.7–0.9, and at least 64 rays provide reasonable initial ranges for evaluation. These values should not be regarded as universally optimal, but rather as a practical calibration space from which system-specific refinement can proceed through iterative inspection of the resulting cavity geometry. Importantly, dynamic binding pockets are inherently heterogeneous, and no single parameterization will perfectly describe every conformation sampled across a structural ensemble.

Beyond quantitative comparison of conformational ensembles, robust pocket characterization has broader applications in structure-based drug discovery and protein engineering. Three-dimensional maps of the accessible pocket volume identify regions that remain consistently available for ligand occupation, providing guidance for where chemical elaboration of a lead compound may be accommodated or where steric clashes are likely to occur as the pocket fluctuates. Likewise, visualization of the molecular surface enclosing the cavity can identify residues that constrain pocket geometry and therefore represent attractive targets for mutagenesis aimed at altering ligand affinity or specificity. Because the methodology described here is computationally efficient and readily applicable to large structural ensembles, it provides a foundation for relating conformational landscapes to molecular recognition, allostery, and rational ligand design.

As increasingly large experimental and computational structural ensembles become available, reproducible methods for characterization of dynamic binding pockets will become increasingly important. The iterative approach presented here provides practical guidance for developing robust and reproducible *measure volinterior* parameterizations while establishing a standardized methodology for quantitative comparison of pocket volume, geometry, and topology across diverse protein systems. By emphasizing internally consistent parameterization across conformational ensembles rather than optimization for individual structures, this approach enables differences in pocket geometry to be interpreted as consequences of protein conformational dynamics rather than artifacts of cavity definition.

## Acknowledgments

This work was funded by the National Institutes of Health (NIH) through award P20GM104316-10 to J.A.H.-P and the University of Delaware. Computer time for MD trajectory analysis on Delta at the National Center for Supercomputing Applications at the University of Illinois at Urbana-Champaign and Anvil at the Rosen Center for Advanced Computing at Purdue University was provided by allocation BIO-240029 from the Advanced Cyberinfrastructure Coordination Ecosystem: Services & Support (ACCESS) program, funded by NSF awards #2138259, #2138286, #2138307, #2137603, and #2138296. This research was supported by the Delaware Advanced Research Workforce and Innovation Network (DARWIN, NSF award OAC-1919839) and the BioStore resource made possible by the National Institutes of Health (NIH, awards P20-GM-103446 and S10-OD-028725).

## Supporting Information

### Code Snippet S1: Tcl script to generate a dummy-atom lattice to visualize pocket volumes as point clouds.

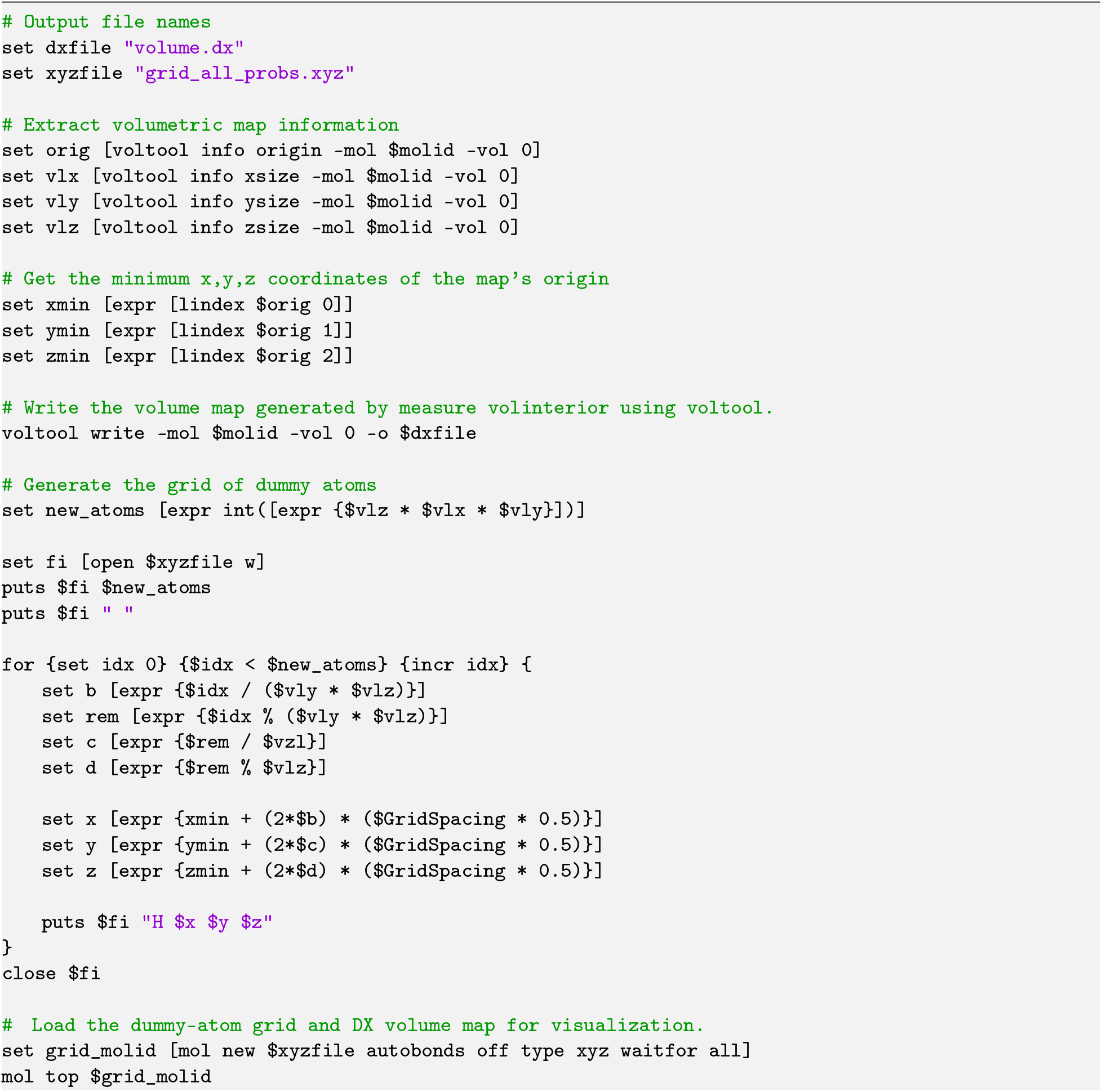
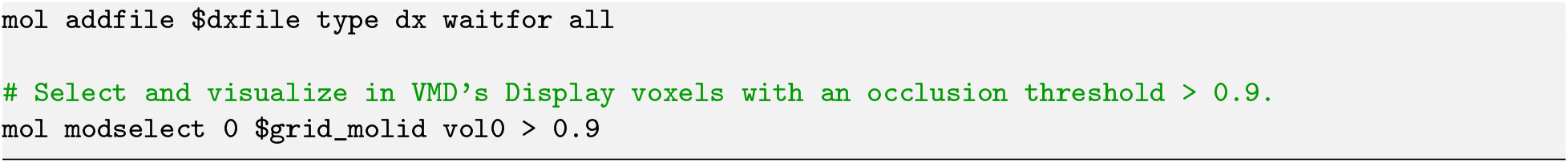

### Code Snippet S2: Tcl script to calculate CAM binding site volume based on a ligand-guided pocket definition.

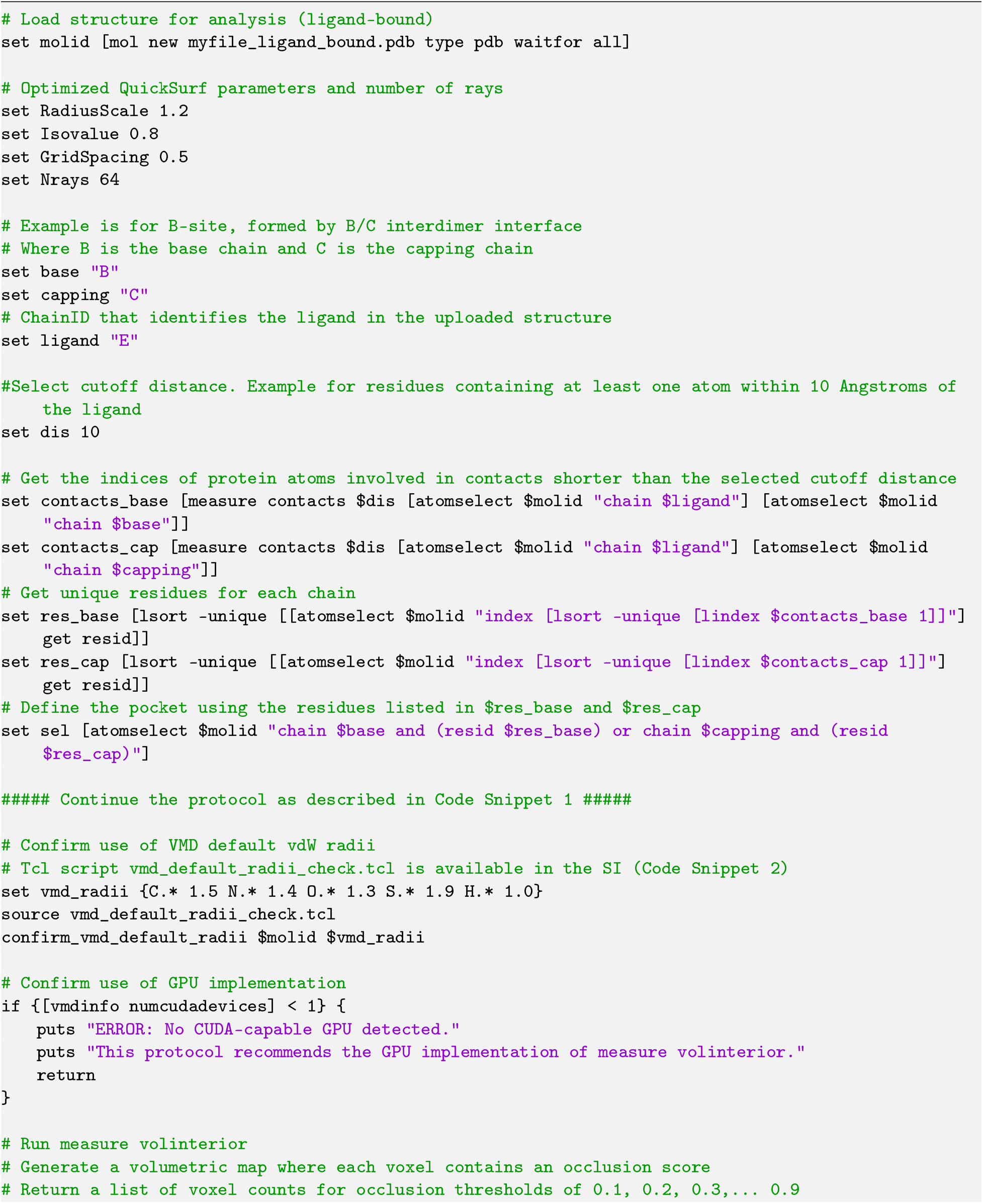
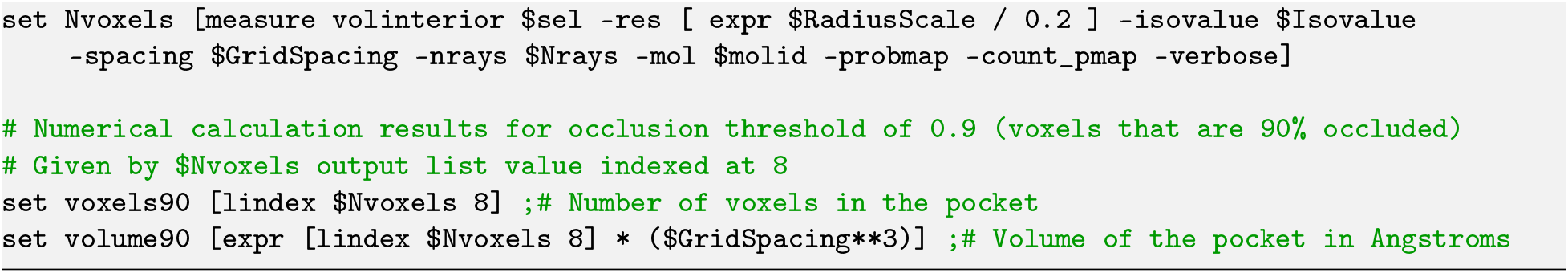

### Code Snippet S3: Tcl script to calculate relative SASA for HBV Cp interdimer interfaces.

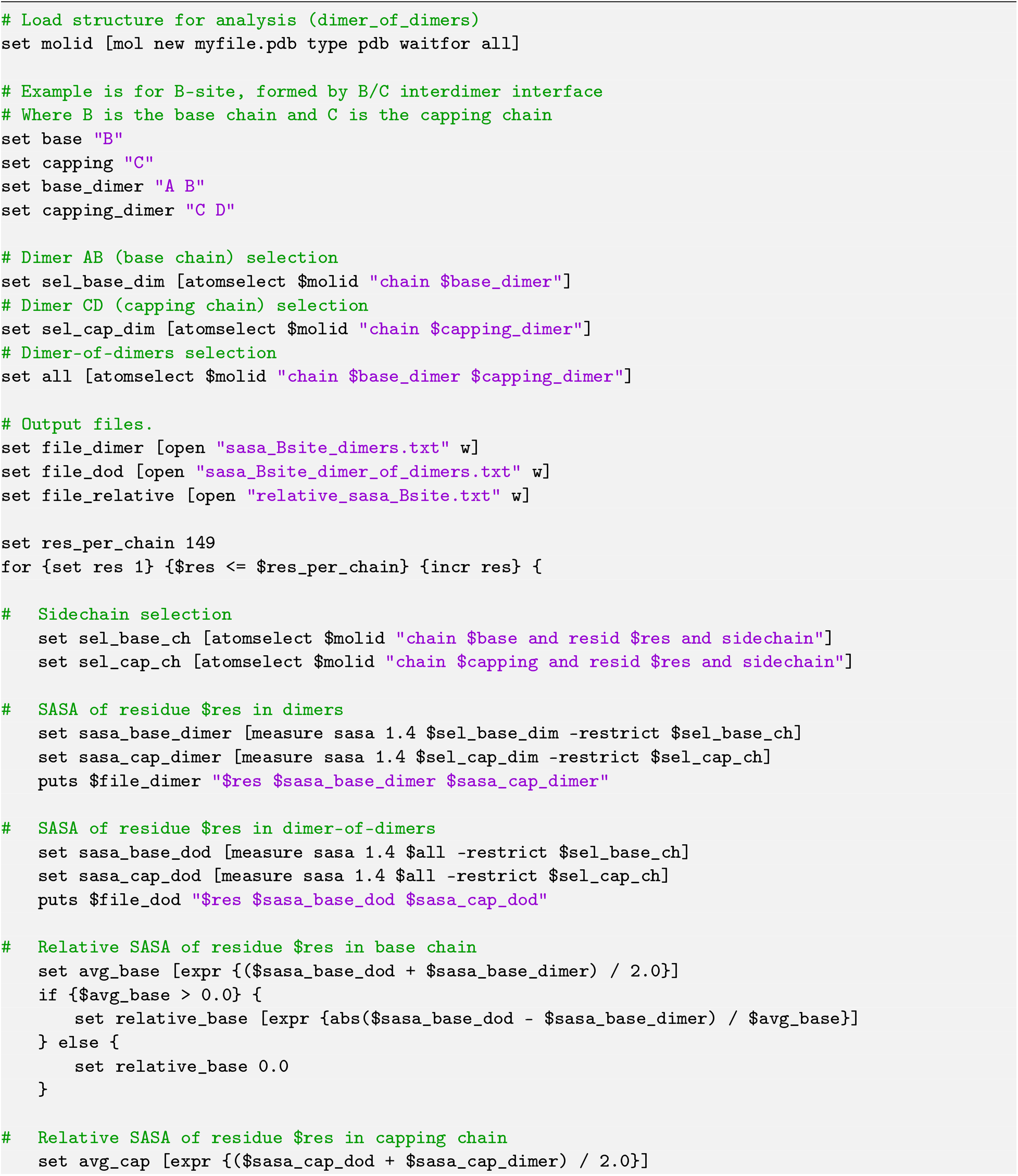
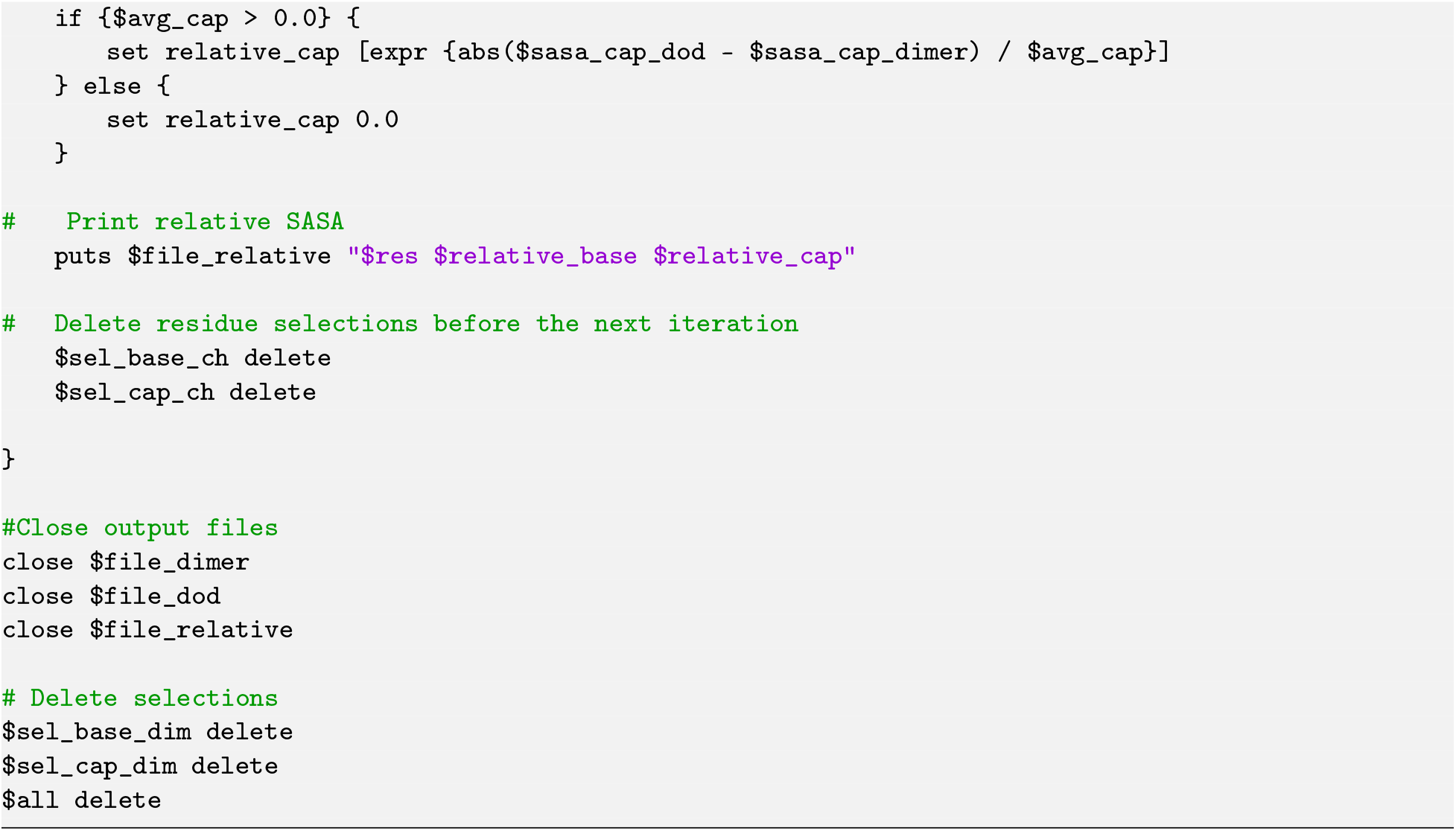

### Code Snippet S4: Tcl script to confirm VMD default vdW radii are set, and otherwise reassign them.

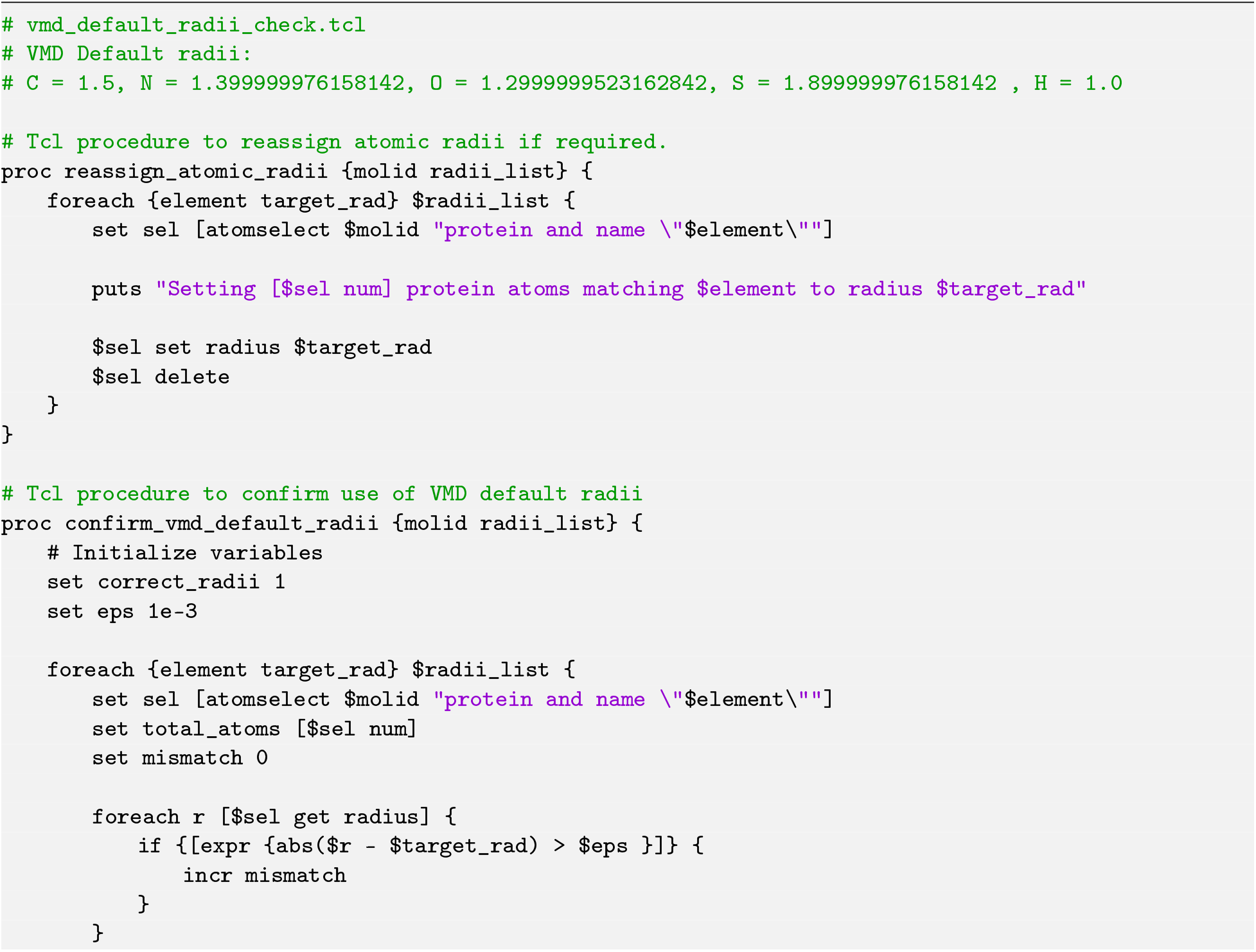
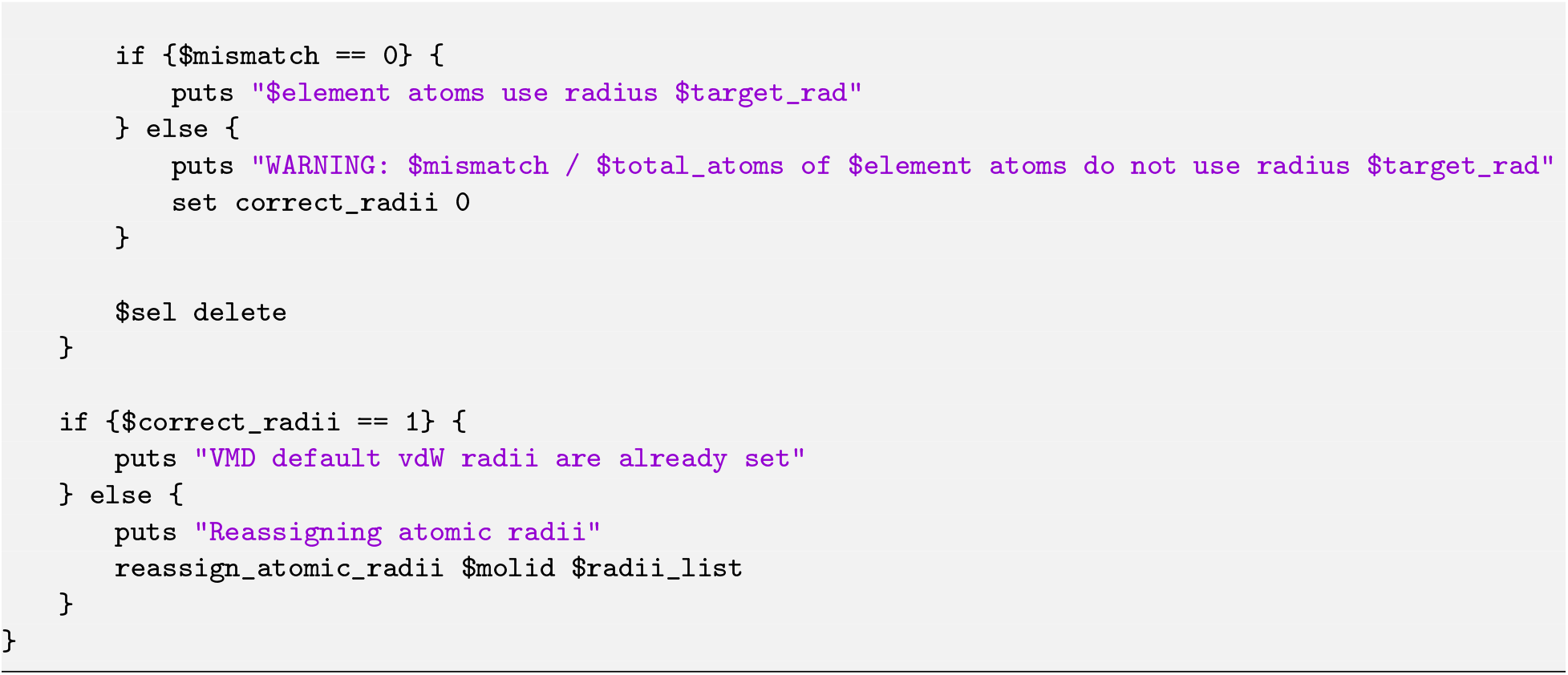

**Figure S1:**
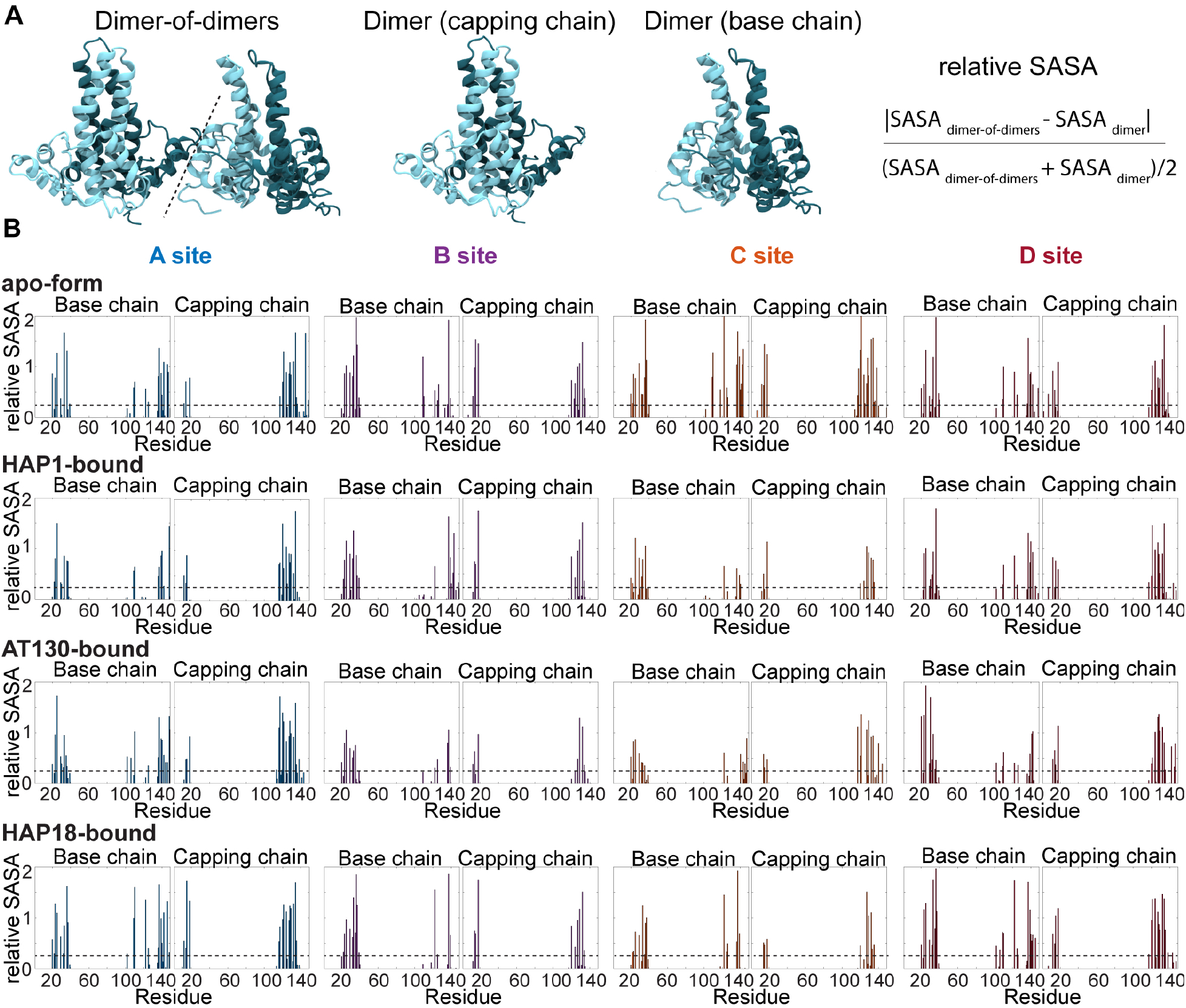
Using relative solvent-accessible surface area to identify interface-forming residues. (**A**) Definition of relative SASA used to identify residues participating in formation of the CAM-binding interface. For each residue, side-chain SASA was calculated for the isolated homodimers of the base and capping chains (*SASA*_dimer_) and for the corresponding dimer-of-dimers surrounding the CAM site (*SASA*_dimer-of-dimers_). Relative SASA was calculated as the absolute difference between these values normalized by their average [57]. (**B**) Relative SASA values for residues in the base and capping chains of the four quasi-equivalent CAM-binding sites across the analyzed crystal structures. The dashed line indicates the relative SASA threshold of 0.25 used to classify interface-forming residues. Residues exhibiting little change in solvent accessibility upon formation of the dimer-of-dimers remain near zero, whereas residues that become buried at the interdimer interface exhibit progressively larger relative SASA values.

**Figure S2:**
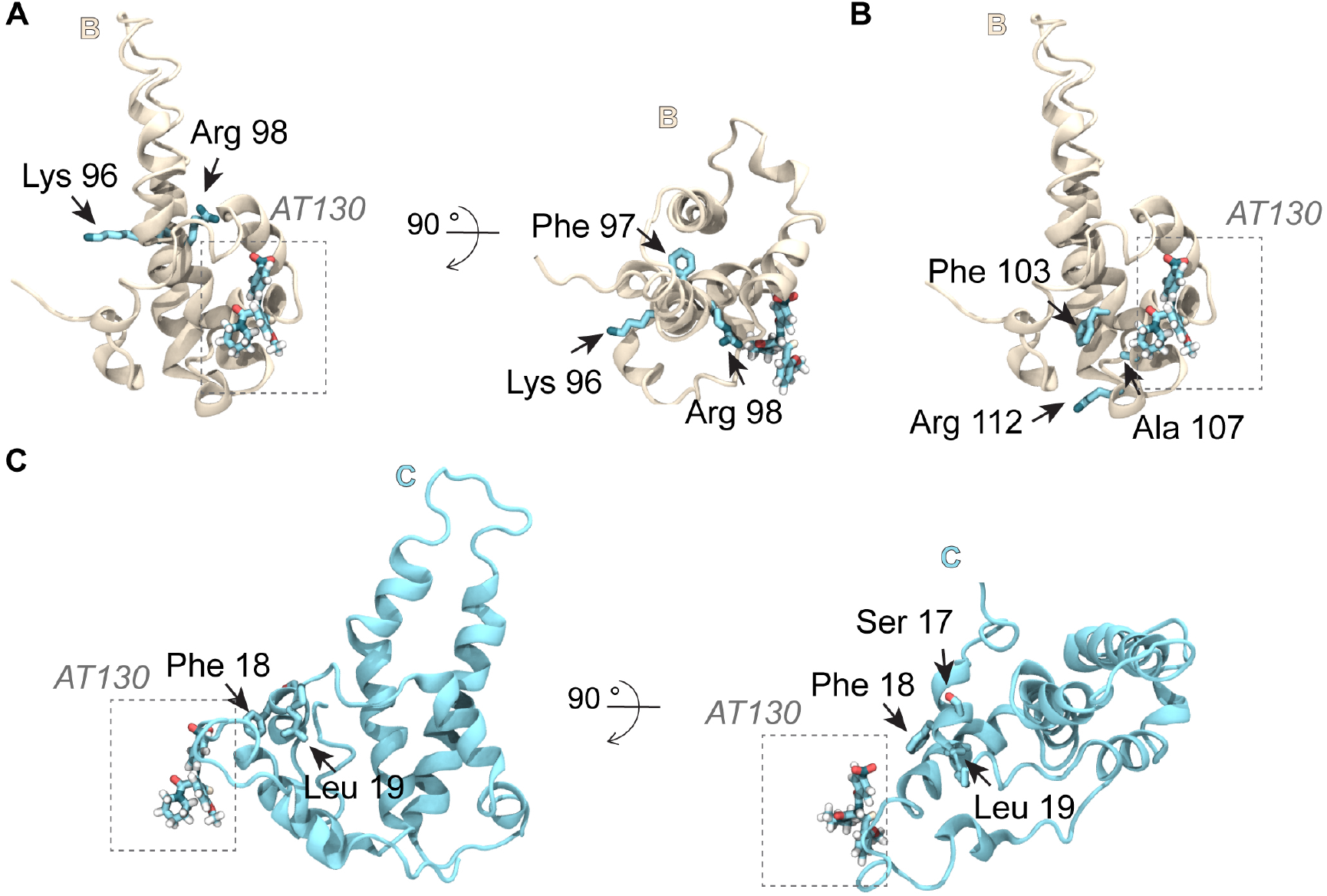
Side-chain orientation provides a final refinement criterion for localizing the molecular container. (**A**) Side-chain orientations of Lys96, Phe97, and Arg98 in the base chain. Arg98 projects toward the fulcrum and contributes to local enclosure of the CAM pocket, whereas Lys96 and Phe97 project away from the cavity and do not contribute to its boundary. (**B**) Side-chain orientations of fulcrum residues in the capping chain. Phe18 projects toward the CAM pocket and improves local enclosure of the cavity, whereas neighboring residues such as Ser17 and Leu19 project away from the binding site and would contribute enclosure of space outside the CAM pocket. (**C**) Side-chain orientations of selected helix 4b residues in the base chain. Phe103, Ala107, and Arg112 project away from the CAM pocket and were therefore excluded from the molecular container because they contribute to extraneous cavity components rather than the binding site itself.

